# Gene expression plasticity and frontloading promote thermotolerance in *Pocillopora* corals

**DOI:** 10.1101/398602

**Authors:** K. Brener-Raffalli, J. Vidal-Dupiol, M. Adjeroud, O. Rey, P. Romans, F. Bonhomme, M. Pratlong, A. Haguenauer, R. Pillot, L. Feuillassier, M. Claereboudt, H. Magalon, P. Gélin, P. Pontarotti, D. Aurelle, G. Mitta, E. Toulza

## Abstract

Ecosystems worldwide are suffering from climate change. Coral reef ecosystems are globally threatened by increasing sea surface temperatures. However, gene expression plasticity provides the potential for organisms to respond rapidly and effectively to environmental changes, and would be favored in variable environments. In this study, we investigated the thermal stress response in *Pocillopora* coral colonies from two contrasting environments by exposing them to heat stress. We compared the physiological state, bacterial and Symbionaceae communities (using 16S and ITS2 metabarcoding), and gene expression levels (using RNA-Seq) between control conditions and heat stress (the temperature just below the first signs of compromised health). Colonies from both thermal regimes remained apparently normal and presented open and colored polyps during heat stress, with no change in bacterial and Symbionaceae community composition. In contrast, they differed in their transcriptomic responses. The colonies from Oman displayed a more plastic transcriptome, but some genes had a higher basal expression level (frontloading) compared to the less thermotolerant colonies from New Caledonia. In terms of biological functions, we observed an increase in the expression of stress response genes (including induction of tumor necrosis factor receptors, heat shock proteins, and detoxification of reactive oxygen species), together with a decrease in the expression of genes involved in morpho-anatomical functions. Gene regulation (transcription factors, mobile elements, histone modifications and DNA methylation) appeared to be overrepresented in the Oman colonies, indicating possible epigenetic regulation. These results show that transcriptomic plasticity and frontloading can be co-occurring processes in corals confronted to highly variable thermal regimes.

## Introduction

Earth is undergoing unprecedented global environmental changes with major effects on biodiversity (Barnosky *et al*. 2011). The ongoing erosion of the most vulnerable ecosystems due to current environmental degradation is particularly worrying and is only a premise to what scientists have called the sixth mass extinction (Barnosky *et al*. 2011). In particular, climate change, ocean acidification and extreme climatic events have already resulted in the irreversible degradation of more than 20% of coral reefs worldwide (Bellwood *et al*. 2004; Hoegh-Guldberg *et al*. 2007). Scleractinian corals constitute the biological and physical framework for a large diversity of marine organisms [c.a. ∼600 coral, ∼2000 fish, and ∼5000 mollusk species (Veron & Stafford-Smith 2000; Reaka-Kudla 2005)]. Hence, the extinction or even major decrease of corals would have dramatic repercussions on the overall associated communities (Hughes *et al*. 2017a). Natural variation in thermal tolerance exists among coral populations (Oliver & Palumbi 2010; Palumbi *et al*. 2014), especially along a latitudinal gradient (Polato *et al*. 2010; Dixon *et al*. 2015), hence providing some hope for coral survival based on their capacity to cope with heat stress. More specifically, it has been shown that populations inhabiting in zones with more variable temperature regimes display better tolerance to heat stress from local (Kenkel *et al*. 2013) to geographical scales (Hughes *et al*. 2003; Riegl *et al*. 2011; Coles & Riegl 2013).

It nevertheless remains unclear whether thermo-tolerance is acquired via acclimation (i.e. intra-generational gene expression plasticity (Barnosky *et al*. 2011; Kenkel & Matz 2016)) and/or through genetic adaptation (i.e. inter-generational microevolution (Barnosky *et al*. 2011; Dixon *et al*. 2015)). Actually, some studies strongly suggest that both processes are likely to co-occur in wild coral populations (Hoegh-Guldberg *et al*. 2007; Reusch 2013; Palumbi *et al*. 2014; Hughes *et al*. 2017a; Torda *et al*. 2017).

With the recent advances of high throughput molecular methods, it is now possible to provide a more precise account of the molecular mechanisms underlying coral response to heat stress. In particular, recent studies clearly demonstrated that coral responses to heat stress involve the fine-tuned regulation of expression levels of some genes/proteins involved in several molecular pathways such as metabolism, stress-response and apoptosis (Brown *et al*. 2002; Weis 2008; Ainsworth *et al*. 2011; Barnosky *et al*. 2011; Bellantuono *et al*. 2012a; Barshis *et al*. 2013; Kenkel *et al*. 2013; Palumbi *et al*. 2014). In this regard, two main molecular patterns having different temporalities have been put forward: (1) “transcriptional plasticity”, i.e. extensive changes in gene expression levels according to the occurring thermal condition and (2) “transcriptional frontloading”, i.e. the elevation of stress related genes baseline expression that preconditions organisms to subsequent (recurrent) stresses (Reaka-Kudla 2005; Mayfield *et al*. 2011; Barnosky *et al*. 2011; Barshis *et al*. 2013; Palumbi *et al*. 2014; Hughes *et al*. 2017a). While such elevated constitutive gene expression levels could reflect local adaptation (i.e. genetically fixed gene expression level (Bellwood *et al*. 2004; Hoegh-Guldberg *et al*. 2007; Oliver & Palumbi 2010; Palumbi *et al*. 2014), it could also reflect an acclimation via epigenetic processes leading to constitutive gene expression (Veron & Stafford-Smith 2000; Reaka-Kudla 2005; Torda *et al*. 2017). Epigenetic changes through environmental priming (i.e. translation of environmental cues) may be involved in adaptive evolution at such short timescales, eventually enabling transgenerational plasticity (Hughes *et al*. 2017a; Torda *et al*. 2017; Jablonka 2017).

Surprisingly, frontloading and gene expression plasticity were generally discussed as mutually exclusive patterns (Oliver & Palumbi 2010; Barshis *et al*. 2013; Palumbi *et al*. 2014; Dixon *et al*. 2015; Kenkel & Matz 2016) although these two molecular processes most likely co-occur during coral responses to heat stress. In particular, one might expect that the regulation strategy of genes (plasticity versus frontloading) will greatly depend on the molecular pathways in which they are involved and the energetic, physiological, and ultimately fitness cost associated with gene expression. So far, frontloading has been detected for stress response genes such as Heat Shock Proteins (HSPs), apoptosis and tumour suppression factors in resilient coral populations under experimentally simulated heat stress inducing bleaching in the common reef-building coral *Acropora hyacinthus* (Polato *et al*. 2010; Barshis *et al*. 2013; Dixon *et al*. 2015; Kenkel & Matz 2016) and for metabolic genes in populations pre-exposed to warm temperatures in response to long-term heat stress in *Porites astreoides* (Kenkel *et al*. 2013; Palumbi *et al*. 2014). Conversely, in the latter species, plasticity was observed in the expression of environmental stress response genes (Hughes *et al*. 2003; Riegl *et al*. 2011; Coles & Riegl 2013; Kenkel & Matz 2016), hence challenging the patterns observed in *A. hyacinthus* (Barnosky *et al*. 2011; Barshis *et al*. 2013; Coles & Riegl 2013; Kenkel & Matz 2016). Although both strategies (i.e. constitutive frontloading *versus* expression plasticity) undoubtedly exist in wild coral populations, the pre-exposure conditions that foster their induction and their relative effects on coral resistance to heat stress still remain unclear (but see (Hughes *et al*. 2003; Barnosky *et al*. 2011; Dixon *et al*. 2015; Kenkel & Matz 2016)).

Importantly, scleractinian corals are composed of several symbiotic organisms including the cnidarian host, the mutualist photosynthetic algae (formerly defined as belonging to the genus *Symbiodinium* but now considered as different genera within the family Symbionaceae (Bellwood *et al*. 2004; Barnosky *et al*. 2011; Dixon *et al*. 2015; LaJeunesse *et al*. 2018)) and bacterial communities. All symbionts involved in a stable symbiosis effectively form the entire organism, and constitute what is referred to the holobiont (Margulis & Fester 1991; Hoegh-Guldberg *et al*. 2007; Reusch 2013; Palumbi *et al*. 2014; Torda *et al*. 2017). A decade after this term was defined, its use has been particularly popularized in reference to corals (Rohwer *et al*. 2002), and subsequent research has led to the hologenome theory of evolution (Rosenberg *et al*. 2007; Zilber-Rosenberg & Rosenberg 2008). In this context, the hologenome is defıned as the sum of the genetic information of the host and its symbiotic microorganisms. Phenotypes are thus the product of the collective genomes of the holobiont partners in interaction with the environment, which constitute the unit of biological organization and thus the object of natural selection (Zilber-Rosenberg & Rosenberg 2008; Guerrero *et al*. 2013; McFall-Ngai *et al*. 2013; Bordenstein & Theis 2015; Theis *et al*. 2016). Additionally to the cnidarian host response, the genotype -or association of genotypes-of the photosynthetic mutualist Symbionaceae symbionts plays a key role in the thermotolerance of the holobiont (Hume *et al*. 2013; Mayfield *et al*. 2014; Suggett *et al*. 2017). There is less certainty about the importance of the coral bacterial community in participating to the fitness of the holobiont, although accruing evidences strongly suggest their implication in coral response to environmental conditions (Li *et al*. 2014; Pantos *et al*. 2015; Hernandez-Agreda *et al*. 2016), and in the resistance to diseases (Sato *et al*. 2009; Cróquer *et al*. 2013; Meyer *et al*. 2016). Finally, the role of the coral-associated microorganisms and their potential to modify holobiont response to stress remain so far overlooked (but see (Ziegler *et al*. 2017; Torda *et al*. 2017). Hence, studying how corals respond to stress implies an integrative approach to analyze the response of each symbiotic protagonist.

With this aim, we investigated the molecular mechanisms underlying thermo-tolerance of coral holobionts. We analyzed the holobiont response to stress of two coral populations originating from environments with contrasting thermal regimes. We used scleractinian corals from the genus *Pocillopora* as model species because they have abroad spatial distribution throughout the Indo-Pacific (Veron & Stafford-Smith 2000). The genus *Pocillopora* is considered to be one of the most environmentally sensitive (van Woesik *et al*. 2011) but its widespread distribution clearly suggests potential for acclimation and/or adaptation which may be correlated to specific differences (i.e. different cryptic lineages may be adapted to different environmental conditions). In particular, we focused on *Pocillopora damicornis-like* colonies from two localities with contrasting thermal regimes: colonies from New Caledonia (NC) are exposed to temperate and stable temperatures over the year, while those from Oman are exposed to globally warmer and more seasonal fluctuating temperatures. As the *corallum* macromorphology is not a discriminant character in *Pocillopora* and as the taxonomic revision of this genus using molecular data reveals that some of the *Pocillopora* species (Schmidt-Roach *et al*. 2014; Gélin *et al*. 2017b) are actually species complexes, we identified *a posteriori* the species of the sampled colonies (mitochondrial ORF sequencing and individual clustering) in order to interpret the results in a precise evolutionary context. To avoid biases inherent in transplantation-based field experiments resulting from environmental factors other than temperature, we undertook our comparative study in a controlled environment in which we mimicked ecologically realistic heat stress to compare the responses of colonies from both localities. We combined a specific RNA-seq approach to study the cnidarian host response, and metabarcoding analyses using ITS-2 and 16S amplicon sequencing to study the dynamics of the associated algal (Symbionaceae) and bacterial community compositions, respectively. According to the literature we first expected to detect changes in both symbiotic algal and bacterial communities in corals from both localities when exposed to heat stress. Moreover, since variable environments are expected to select for plasticity, we predicted that the cnidarian hosts from Oman may display more gene expression plasticity than those from New Caledonia. However, because frontloading was also found to be an alternative response to recurrent changing conditions, we might also expect some degrees of constitutive high levels of gene expression at least for some molecular pathways and more particularly in Oman corals.

## Material and Methods

### Coral sampling and maintenance

*Pocillopora damicornis*-like colonies originating from environments characterized by contrasting thermal regimes were sampled during the warmer month in two different localities: (1) in Oman, Gulf of Oman, Northwestern Indian Ocean (Om; June 2014; local seawater temperature during sampling 30.8°C), where corals are exposed to a globally warmer and variable thermal environment, and (2) in New Caledonia, Southwestern Pacific Ocean (NC; November 2014; local seawater temperature during sampling 27.1°C), where corals are subject to more mitigate and stable temperatures (see Table 1 for the sampling sites and Table 2 for temperature regime of locality).

**Table 1.**
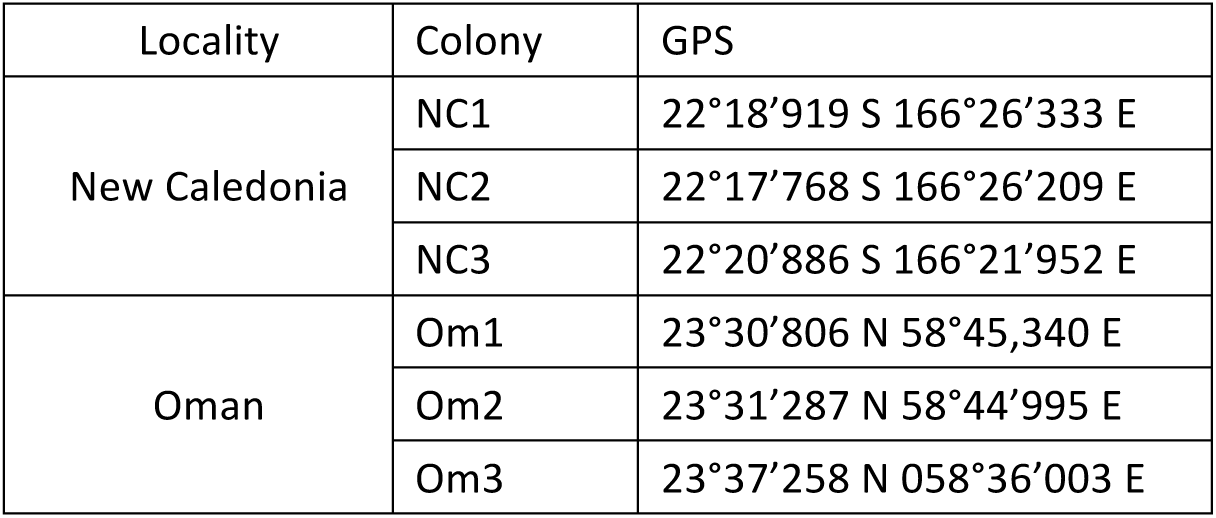
Sampling sites

**Table 2.** Sea Surface Temperatures (SST) to which the colonies sampled in this study are exposed in their natural environments. Thermal regime descriptors were compiled from weekly mean sea surface temperature data collected from the Integrated Global Ocean Services System Products Bulletin (IGOSS: http://iridl.ldeo.columbia.edu/SOURCES/.IGOSS/) for quadrats of 1° longitude x 1° latitude from 1982 to the year of sampling (2014).

|  | New Caledonia | Oman |
| --- | --- | --- |
| Mean SST (°C) | 24.8 | 27.9 |
| Variance (°C) | 2.7 | 9.5 |
| Min SST (°C) | 22.6 | 22.1 |
| Max SST (°C) | 27.1 | 33.2 |
| Mean SST of 3 warmer months (°C) | 26.8 | 31.3 |
| Mean SST of 3 cooler months (°C) | 22.8 | 23.8 |

From each location, we thus sampled colonies morphologically similar and occupying the same water depth niche. To account for possible intra-population diversity, three colonies (>20 cm in diameter) were collected in each locality, and separated by at least 10 m to decrease the probability to collect members of the same genet, as some *Pocillopora* species are able to propagate by asexual reproduction (Adjeroud *et al*. 2013; Gélin *et al*. 2017a; 2018).

Immediately following collection, a 1 cm branch tip of each colony was excised, rinsed three times in filtered seawater (0.22 µm), and placed in RNAlater solution (Sigma Aldrich) for the *in situ* microbiota analysis. The rest of the colony was fragmented into 20 branches each of 10 cm length and physiologically stabilized in openwater system for one week before shipping (Al-Hail field stationof the Sultan Qaboos University and the Public aquarium of Noumea for OM and NC localities respectively). For shipping, individual branches were placed in plastic bags containing oxygenated seawater (800mL seawater and 1600mL of medical oxygen), and transported by aircraft to the research aquarium of the Banyuls-sur-Mer oceanographic observatory (France). The coral branches were maintained in artificial seawater (Seachem Reef Salt) at 26°C, and supplied daily with *Artemia* nauplii to satisfy their heterotrophic demand. The conditions in the maintenance tank were controlled to mimic the natural physicochemical parameters of coral reefs (pH:8.2; salinity: 36 psu; light intensity: 150 to 250 µmol of photons/m²/s; photoperiod: 12h night/12h day;kH: 6–7.5 dkH; calcium concentration: 410–450 mg/L; inorganic phosphate concentration: < 0.1 mg/L; magnesium concentration: 1300–1400 mg/L; nitrate concentration: < 5 mg/L). After 3 and 7 months of acclimatization to the laboratory condition (marked by growth resumption) for Om and NC colonies, respectively, corals were fragmented to produce a total of ∼15 to 20 clones (nubbins) from each colony (∼3 cm). These were individually fixed to a support (here a p1000 tip) using an epoxy adhesive. We waited for complete healing (evident as tissue extending to cover the epoxy adhesive) prior to run the experiment.

### Ecologically realistic heat stress

The aim of this experiment was to compare the response to heat stress of colonies from two localities having the same physiological state, to investigate the patterns of expression of the molecular pathways involved during the stress exposure and the putative modifications of the coral microbiota.

The experimental design comprised four tanks of 53 L per locality in which the seawater was continuously recycled. The water was sterilized using UV (rate 3200 L/h) and renewed twice per hour in each tank (recirculation rate: 100L/h in each tank). The eight tanks shared the same seawater but their temperature was monitored individually (HOBBY BiothermPro, model 10892; 500W Aqua Medic titanium heater; HOBO TidbiT v2 logger) (Supplementary Figure S1). For each locality, 5 to 8 nubbins per mother colonies were randomly placed in each tank (four tanks per locality) for two weeks at the control temperature and the following protocol was applied: three tanks were then subjected to a gradual temperature increase (stress treatment) while the fourth (control) was maintained at the control temperature to verify that the stress observed in the stressful treatment was not due to other potential confounding effects or water cues (Figure 1). Both the control and stress temperatures were specific for each sampling locality to mimic their respective natural environment. In particular, we set the control temperature as the mean water temperature for the three warmer months measured at the coral sampling site locality (Table 1): 31°C for the colonies from Om, and 27°C for the colonies from NC. The stress treatment was ecologically realistic, i.e. reflecting a naturally occurring warming anomaly, and consisted in increasing the temperature gradually by 1°C (over 5 consecutive hours) each week until physiological collapse of the corals became evident (polyps closure, bleaching or necrosis), as described by (Vidal-Dupiol *et al*. 2009). Sampling was performed in the three sampling tanks just before the first temperature increase (control condition) as well as each week before the next temperature increase. The beginning of polyp closure was consistently observed for the different colonies of the same locality at the same temperature threshold. Samples for subsequent genetic and transcriptomic analyses were chosen *a posteriori*. They corresponded to those sampled in each tank just before the first increase of temperature (control samples), and just before the temperature that produced the first signs of physiological collapse and before bleaching (stress temperature samples). Thus, for each condition (control and stress) we obtained three biological replicates of each colony from the three different tanks (three colonies per locality) to reach a total of 36 samples (2 localities × 3 colonies× 2 experimental conditions × 3 replicates/tanks). The general health of the nubbins was assessed via daily photographic monitoring (at noon prior to feeding) throughout the period of the experiment.

**Figure 1.**
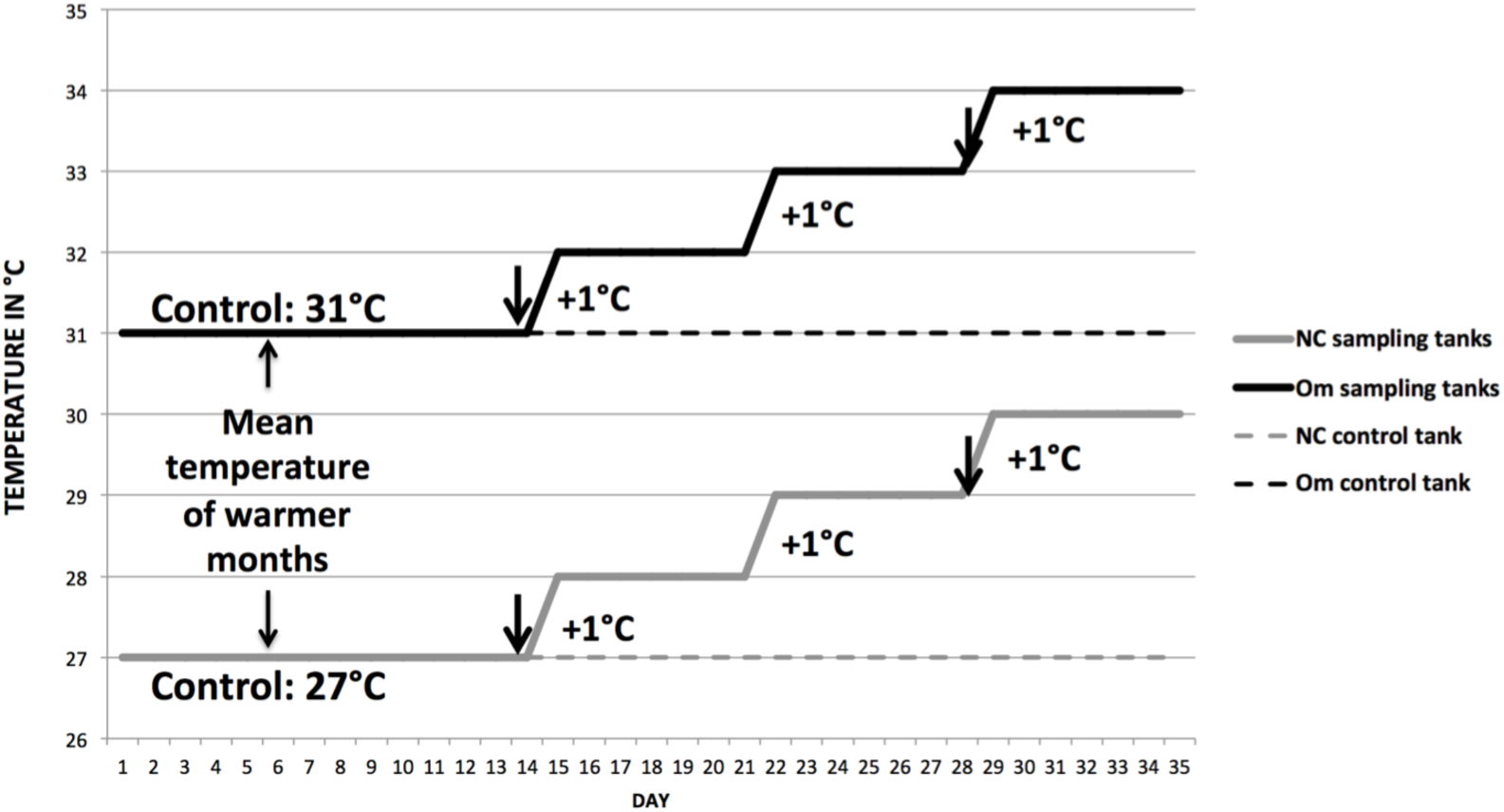
The ecologically realistic heat stress experiment: from mean temperatures of the warmer months in natura to a pre-bleaching physiological state. Nubbins were collected at each time point and arrows represent points at which nubbins were chosen for analyzing the microbial composition and the transcriptomic response of the host.

### DNA extraction

DNA was extracted from each 36 samples as well as coral tips directly collected on the six colonies *in natura* for the *in situ* condition (three in Om, three in NC), using the DNeasy Blood and Tissue kit (Qiagen) following the manufacturer’s instructions. DNA was quantified by spectrophotometry (NanoDrop).

### Host species and clonemates identification

As the *corallum* macromorphology is not a diagnostic criterion in *Pocillopora* genus, the host species was thus identified molecularly. Thus each colony was sequenced for the mitochondrial variable open reading frame (ORF) and was genotyped using 13 specific microsatellites, as in Gélin *et al*. (Gélin *et al*. 2017b). Then each colony used in the experiment was assigned to Primary and Secondary Species Hypothesis (PSH and SSH; *sensu* Pante et al.) (Pante *et al*. 2015) following the nomenclature from Gélin *et al*. (Gélin *et al*. 2017b). Indeed, sampling *Pocillopora* colonies presenting various morphs from different locations from the Indo-Pacific, Gélin et al classified these colonies, without *a priori* based on *corallum* macromorphology, into Species Hypotheses (*sensu* Pante et al., i.e. the species are hypotheses that can be confirmed or refuted while new data are added) (Pante *et al*. 2015) using sequence-based species delimitation methods, a first sorting allowed to define Primary Species Hypotheses (PSH) and then individual clustering based on microsatellite multilocus genotypes allowed a second sorting delimiting Secondary Species Hypotheses (SSH). Thus comparing the ORF sequences obtained in this study to those from (Gélin *et al*. 2017b), the sampled colonies were assigned to a PSH. Then, if relevant, the colonies were assigned to SSH performing clustering analysis using Structure 2.3.4 (Pritchard *et al*. 2000), as in (Gélin *et al*. 2017b). Meanwhile, the identical multi-locus genotypes (i.e. clonemates if any) were identified by microsatellite analysis using GenClone (Arnaud-Haond & Belkhir 2006) as in Gélin *et al*. (Gélin *et al*. 2017a).

### Microbial community analysis using MiSeq 16S and ITS2 metabarcoding

The aim of this analysis was to investigate the composition and the dynamics of the two principal symbiotic coral communities (i.e. bacterial and algal) *in situ* and during heat stress.

### Amplicon Sequencing

A bacterial 16S rDNA amplicon library was generated for each of the 42 samples (one *in situ* condition, three control conditions and three stress conditions per colony, three colonies per locality, two localities), using the 341F (CCTACGGGNGGCWGCAG) and 805R (GACTACHVGGGTATCTAATCC) primers, which target the variable V3/V4 loops (Klindworth *et al*. 2012). The Symbiodiniaceae assemblages were analyzed using ITS2 (internal transcribed spacer of the ribosomal RNA gene) amplicon libraries and specific primers targeting a sequence of approximately 350 bp (ITS2-F GTGAATTGCAGAACTCCGTG; ITS2-R CCTCCGCTTACTTATATGCTT) (Lajeunesse & Trench 2000; Quigley *et al*. 2014). For both markers, paired-end sequencing using a 250 bp read length was performed on the MiSeq system (Illumina) using the v2 chemistry, according to the manufacturer’s protocol at the Centre d’Innovation Génome Québec and McGill University, Montreal, Canada.

### Bioinformatic analysis

The FROGS pipeline (Find Rapidly OTU with Galaxy Solution) implemented on a Galaxy platform (http://sigenae-workbench.toulouse.inra.fr/galaxy/) was used for data processing (Escudié *et al*. 2017). In brief, paired reads were merged using FLASH (Magoč & Salzberg 2011). After cleaning and removal of primer/adapters using cutadapt (Martin 2011), *de novo* clustering was performed using SWARM (Mahé *et al*. 2014). This uses a local clustering threshold with an aggregation distance (d) of 3. Chimeras were removed using VSEARCH (Rognes *et al*. 2016). We filtered the dataset for singletons and performed affiliation using Blast+ against the Silva database (release 128, September 2016) for 16S amplicons (Altschul *et al*. 1990). For ITS2 metabarcoding, the Symbiodiniaceae type was assessed using Blast+ against an in-house database of Symbiodiniaceae reference sequences built from sequences publicly available. An OTU table in standard BIOM format with taxonomic affiliation was produced for subsequent analyses.

For community composition analysis we used the *phyloseq* R package (McMurdie & Holmes 2013) to infer alpha diversity metrics at the OTU level (i.e. Shannon and Chao indexes), and beta diversity (between sample similarity) from the OTU table. Community similarity was assessed by Principal Coordinate Analysis (PCoA) using the Bray-Curtis distance matrices.

To test for possible effect of corals’ locality of origin and for temperature treatments on the two alpha diversity metrics (i.e. Shannon and Chao) we ran a series of four generalized mixed models including these two fixed factors and their interactions as explanatory variables. The four models consisted in considering all factors, the two fixed factors without accounting for their possible interactions, and the two factors independently. In all models, we also accounted for possible effect of genotypes that were considered as random factors nested within locality. All models were run using the lme4 package implemented in R (Bates *et al*. 2015) and considering a Gamma distribution. The best model for each alpha diversity metrics was identified based on the Akaike Information Criterion (AIC (Akaike 1974)). Concerning beta diversity, we performed MANOVAs to compare beta diversity metrics among the groups of samples by sampling locality, genotype or treatment.

Corrections based on multiple testing were performed using FDR (Benjamini & Hochberg 1995). For all analyses, the threshold significance level was set at 0.05.

### Transcriptome analysis

The aim of this analysis was to study the transcriptomes of the sampled colonies in response to heat stress compared with controlled conditions.

### RNA extraction

Total RNA was extracted from each coral sample using TRIzol reagent (Invitrogen), according to the manufacturer’s protocol. The quantity and integrity of the total RNA extracted was checked using an Agilent 2100 Bioanalyzer (Agilent Technologies)(mean RIN =7.5). Paired-end fragment libraries (2 × 100 bp) were constructed and sequenced on an Illumina HiSeq 2000 platform at the Centre d’Innovation Génome Québec at McGill University, Montreal, Canada.

### Bioinformatic analyses

Fastq read files were processed on the Galaxy instance of the IHPE (http://bioinfo.univ-perp.fr) (Giardine *et al*. 2005). Quality control and initial cleaning of the reads were performed using the filter by quality program (version 1.0.0) based on the FASTX-toolkit (Blankenberg *et al*. 2010). Reads having fewer than 90% of bases having a Phred quality score ≤ 26 were discarded (probability of 2.5 incorrect base call out of 1000, and a base call accuracy of 99.75%). Adaptors used for sequencing were removed using the cutadapt program version 1.6 (Martin 2011). All paired-end reads were aligned using RNAstar software under default parameters, with at least 66% of the bases being required to align to the reference, and a maximum of ten mismatches per read (Dobin *et al*. 2013). The *Pocillopora damicornis sensu lato* reference genome used in this study (Vidal-Dupiol *et al*. 2019) consisted of a draft assembly of 25,553 contigs (352 Mb total) and N50 = 171,375 bp. The resulting transcriptome served as the reference for reads mapping, and a GTF annotation file was constructed using cufflink/cuffmerge (Trapnell *et al*. 2010). HTseq was used to produce count files for genes (Anders *et al*. 2015). The DESeq2 package was used to estimate the normalized abundances of the transcripts, and to calculate differential gene expression for samples between the control temperature and the stress temperature for each locality (Love *et al*. 2014), considering the different genotypes (three biological replicates for each genotype) and using default parameters. We next analyzed genes according to their expression patterns among the different colonies and temperature treatments. Genes were clustered manually into six groups according to their differential expression levels: common over-expressed genes, NC-specific over-expressed genes, Om-specific over-expressed genes, common under-expressed genes, NC-specific under-expressed genes, and Om-specific under-expressed genes. Cluster 3.0 (de Hoon *et al*. 2004) and Treeview (Saldanha 2004) were used to build the heatmap.

### Discriminant analysis of principal components (DAPC)

Our aim was to quantify and compare the level of genome-wide transcriptome plasticity between colonies from Om and NC in response to heat stress. To achieve this we performed a discriminant analysis of principal components (DAPC) based on a log-transformed transcript abundance matrix (containing 26,600 genes) obtained from the 36 samples (*i.e.* 9 control and 9 stressed replicates per locality), as described previously (Kenkel & Matz 2016). Specifically, we ran a DAPC analysis using the resulting log2 transformed dataset for the colonies from NC and Om reared in controlled conditions as predefined groups in the *adegenet* package implemented in R (Jombart *et al*. 2010). Two principal components and a single discriminant function were retained. We then predicted the position of stressed colonies from both localities (Om and NC) onto the unique discriminant function of the DAPC.

We next ran a general linear model (GLM) using the DAPC scores as dependent variable, and accounted for the locality of origin (NC and Om), the treatment (control and heat stress), and their interaction as explanatory variables. We also considered the effect of individual colonies nested within localities as random effects in the model. Our final objective was to test for a potential significant effect of the interaction between the locality and the treatment effects, as a proxy of significant differences in the genome-wide gene expression reaction norms (i.e. differences in DAPC scores between the control and the heat stress treatments) between Om and NC colonies.

### GO enrichment of differentially expressed genes

The transcriptome was annotated *de novo* using a translated nucleotide query (blastx (Altschul *et al*. 1990)) against the non-redundant protein sequence database (nr). The 25 best hits were then used to search for gene ontology terms using the Blast2Go program (Conesa *et al*. 2016). To identify the biological functions significantly enriched within up or down-regulated genes, a Gene Ontology (GO) term enrichment analysis was performed. Lists of GO terms belonging to significantly up-regulated and down-regulated genes were compared to the GO terms of the whole expressed gene set using a Fischer exact test and a FDR value of 0.05. We used REVIGO to visualize the enriched biological processes (Supek *et al*. 2011).

## Results

### Host identification

Among the three colonies from New Caledonia, colonies NC2 and NC3 presented haplotype ORF18 and were assigned to Primary Species Hypothesis PSH05 and more precisely to Secondary Species Hypothesis SSH05a (Gélin *et al*. 2017b), corresponding to *P. damicornis* type *β* (Schmidt-Roach *et al*. 2014) or type 5a (Pinzón *et al*. 2013), while colony NC1 presented ORF09 and was assigned to PSH04, *P. damicornis* type *α*, *P. damicornis* or type 4a, respectively. As for colonies from Oman, they all presented ORF34 and were assigned to PSH12 (Gélin *et al*. 2017a) or type 7a (Pinzón *et al*. 2013) (Supplementary Table S2), that is not part of the *P. damicornis sensu lato* species complex. Thus NC colonies are phylogenetically closer from each other than from colonies from Oman. These three PSHs represent three different species.

Furthermore, NC2 and NC3 multi-locus genotypes (MLGs) differed only from one allele over 26 gene copies, and were thus part of the same clonal lineage (genet), i.e. the entity that groups together colonies whose multi-locus genotypes slightly differ due to somatic mutations or scoring errors. All the other colonies presented MLG that differed enough not to be considered as clonemates or members of the same clonal lineage (genet).

### Ecologically realistic heat stress

Our goal was to ensure that our experimental heat stress faithfully reflects a realistic heat stress *in natura*. Following collection from the field, the corals from the different localities were first maintained in the same controlled conditions at 26°C prior to the experiment. During this period no mortality or signs of degradation/stress were observed for any of the coral colonies. Two weeks before the experiment, a first acclimatization to the control temperatures (27°C or 31°C for NC and Om respectively) was performed. During the experimental heat stress (i.e. gradual temperature increase), visual and photographic monitoring clearly indicated that the first sign of coral stress (i.e. the closure of polyps) occurred at day 30 for both sampling localities, corresponding to 30°C and 34°C for the NC and Om colonies, respectively. These temperatures perfectly match the warmest temperature experienced by these colonies in the field (Table 1). No sign of physiological collapse were observed in control corals throughout the experiment indicating that all the other parameters were maintained optimal for coral colonies.

### Bacterial communities

Among the overall 42 samples analyzed, a total of 5,308,761 16S rDNA amplicon sequences were obtained after cleaning and singleton filtering corresponding to 15,211 OTUs. In all samples the class Gammaproteobacteria was dominant (77.7%), particularly the genus *Endozoicomonas* (44.7% of the sequences); this genus is known to be an endosymbiont of numerous scleractinians (Neave *et al*. 2016b) (See Supplementary Figure S3 for complete bacterial composition in each colony and replicate).

Concerning the two alpha diversity metrics (i.e. Shannon and Chao indexes), the best statistical models to explain the overall variance included the coral’s locality of origin only (AIC = 101.9). According to these models, colonies from Oman displayed a higher Shannon index than those from New Caledonia (P = 0.04). Conversely, the effect of locality was not significant for the Chao index (P > 0.05). The PCoA of Bray-Curtis distances for all colonies showed no evident clusters based on the experimental treatments (Figure 2). We observed a loose grouping based on localities and colonies, except for colony NC1, which appeared to have a more specific microbiota composition, as it had a different grouping associated with the first axis, which explained 22.3% of the variance. This could be correlated with the different species hypotheses for NC1 compared to NC2 and NC3 (see above). These observations were confirmed by statistical analyses (MANOVA between localities: *P*=0.003; between genotypes: *P*=0.001; between temperature treatments: *P*=0.761; Supplementary Table S4). Thus, the bacterial composition appeared to be relatively specific to each colony within each locality, but no major shift was observed during the transition from the natural environment to artificial seawater, nor during heat stress exposure.

**Figure 2.**
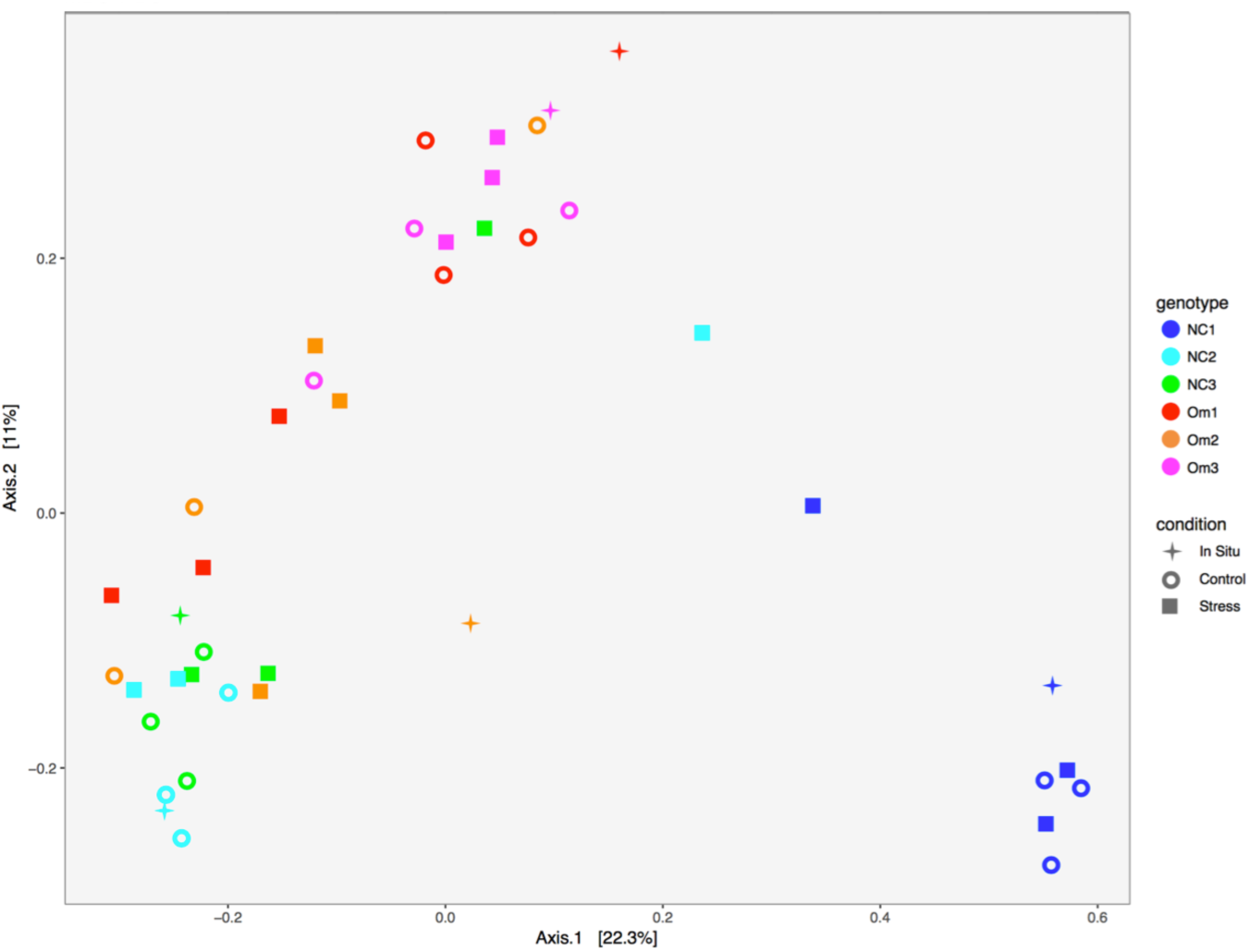
Principal coordinate analysis plot for Bray-Curtis distances of the bacterial composition of each colony in each experimental condition. Different colors represent different colonies, the stars represent the *in situ* conditions, the open circles represent the control conditions, and the squares represent the stress conditions.

### Symbiodiniaceae assemblages

Analysis of the Symbiodiniaceae composition was performed based on an ITS2 metabarcoding, which allowed intra-clade resolution.

Removal of OTUs having an abundance of < 1% left only 4 OTUs among all samples. Two of these corresponded to type C1, while the other two corresponded to type D1a according to (Baker 2003). Type D1a was highly dominant in the colonies originating from Oman, whereas type C1 was almost exclusive to the corals from New Caledonia (Figure 3). The Symbiodiniaceae community composition was very specific to each locality, but remained stable during the transition from the natural environment to artificial seawater, and during heat stress exposure.

**Figure 3.**
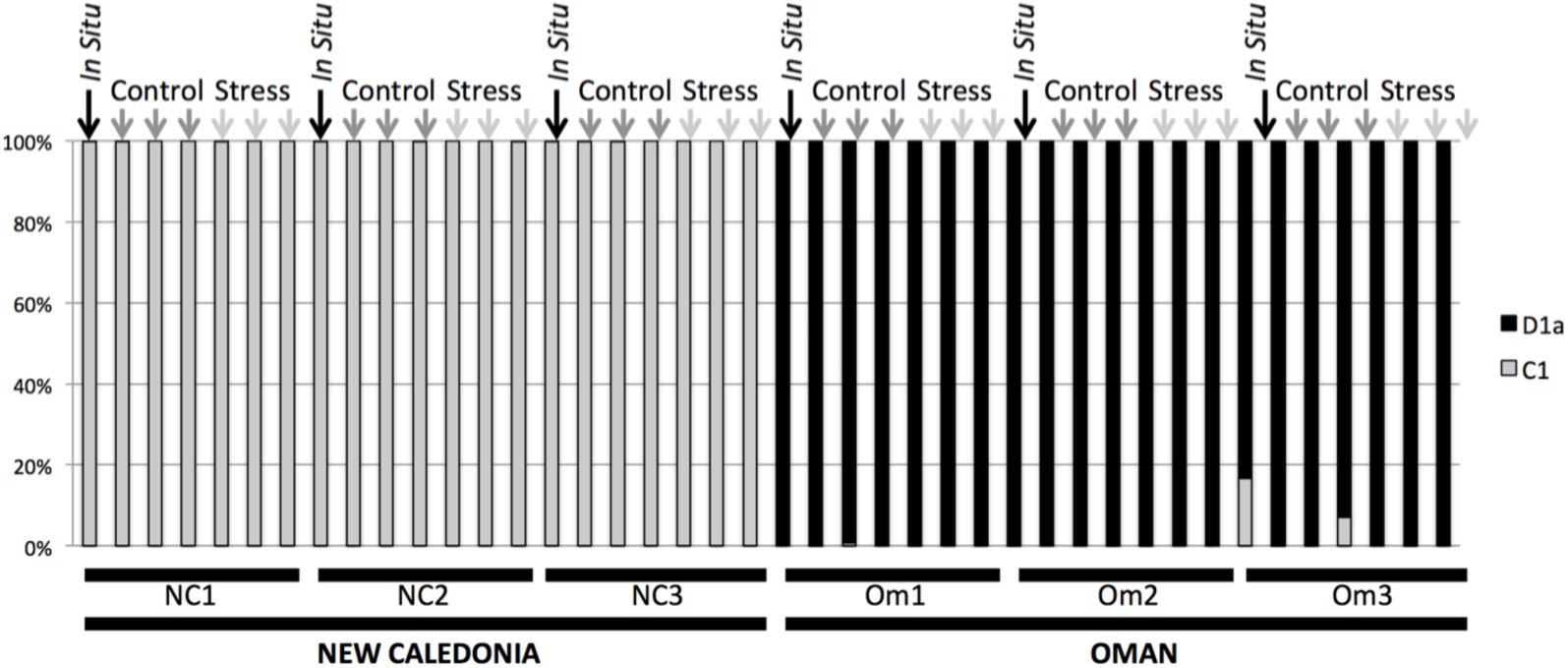
Composition of the Symbiodiniaceae community in each colony in situ and in controlled and stressful experimental conditions.

### Host transcriptome analysis

We generated 36 transcriptomes corresponding to triplicate samples for three colonies of each locality exposed to the control (C) and stress (S) temperatures.

Overall, the transcriptome sequencing of these36 samples yielded 1,970,889,548 high quality Illumina paired reads of 100 bp. Globally, 40–64% of reads obtained for the Om colonies, and 59–70% of reads obtained for NC colonies successfully mapped to the *Pocillopora damicornis* (type β) reference genome. The apparently better alignment of samples from New Caledonia most likely relies on the fact that the New Caledonia colonies used in this study belong to *P. damicornis* types α or 4a (PSH04) and β or 5a (PSH05), which are phylogenetically close to each other and closer from the reference genome, than the Om colonies from type 7a (PSH12) that is phylogenetically more distant from the reference genome. The aligned reads were assembled in 99,571 unique transcripts (TCONS), representing putative splicing variants of 26,600 genes identified as “XLOC” in the genome (FASTA sequences available in Supplementary File S5).

The hierarchical clustering analyses clearly grouped together samples belonging to the same locality and species hypothesis according to their genome-wide gene expression patterns, in link with the phylogenetic differences between the NC and Om haplotypes (Figure 4). Within locality and species hypothesis, the transcriptomes also grouped by colony, indicating that the transcriptomes were genotype-specific. For each colony, the transcriptomes then grouped by condition (control or heat stress), except for New Caledonia colonies NC2 and NC3 (corresponding to the same clonal lineage) that clustered together when exposed to control and heat stress conditions.

**Figure 4.**
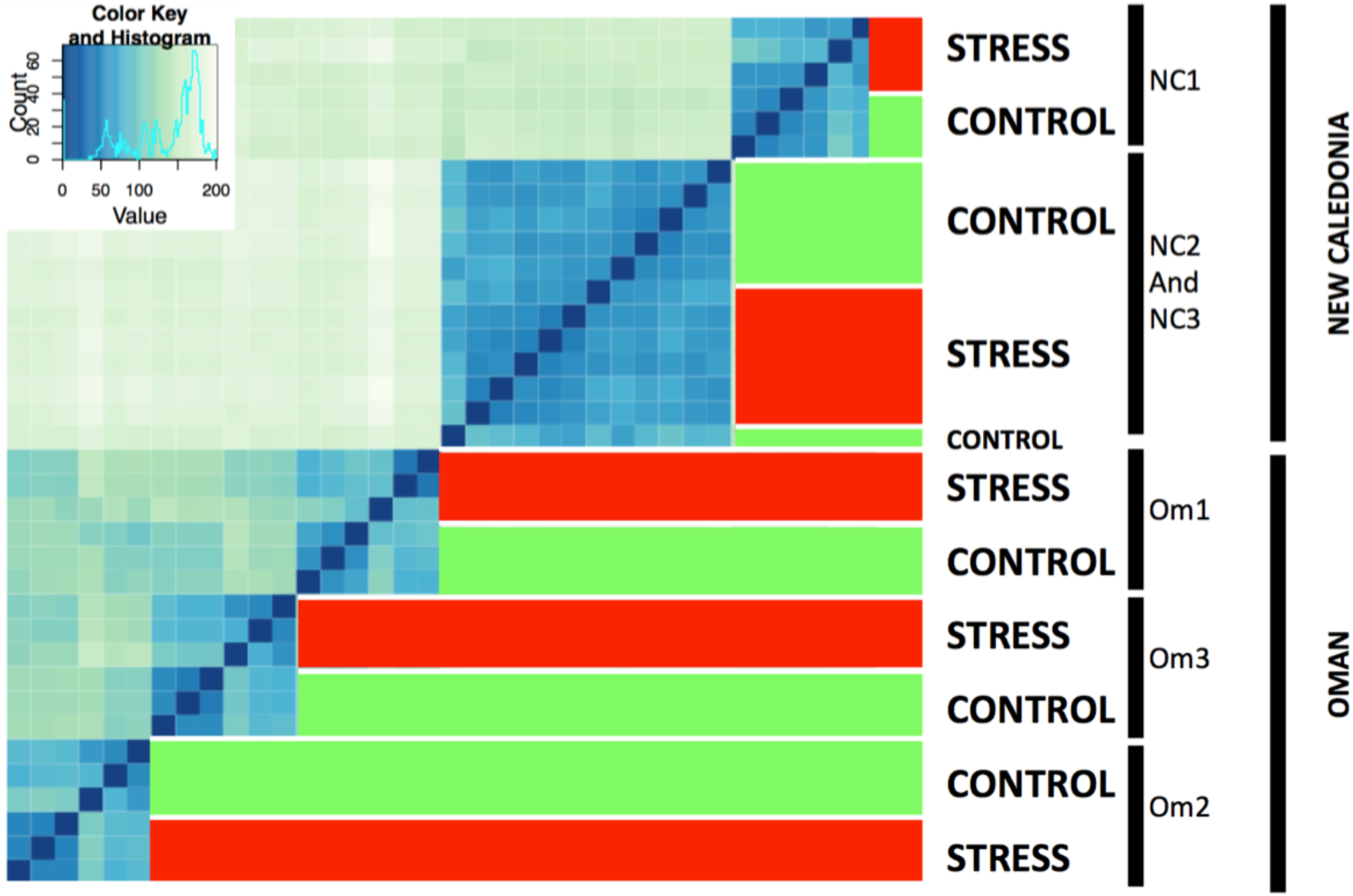
Hierarchical clustering analyses performed using DESeq2 rlog-normalized RNA-seq data for the 36 transcriptomes: two conditions (control and heat stress); three replicates per condition for each colony; three colonies per locality; and two localities [Oman (Om) and New Caledonia (NC)]. The color (from white to dark blue) indicates the distance metric used for clustering (dark blue corresponds to the maximum correlation values).

Despite clustering of the transcriptomes by locality, as the sampling of *Pocillopora damicornis*-like colonies actually corresponded to different species we performed differential gene expression analysis for each colony independently (comparing the biological triplicates for the control condition vs. triplicates for the heat stress conditions). For each locality, the different colonies displayed similar patterns of differential gene expression with in any case a higher number of differentially expressed genes and higher fold-changes between control and heat stress condition in Om compared to NC (Supplementary Figure S6). We detected 673, 479 and 265 differentially expressed genes for NC1, NC2 and NC3 respectively, vs. 2870, 2216 and 959 for Om1, Om2 and Om3. Samples were thus grouped for each locality (nine control nubbins + nine heat stress nubbins) for subsequent analyses (full results of the comparisons between stressed and controls (log2-foldchange and adjusted *p-*values) for each colony or between the two localities are provided in Supplementary File S7). For Om colonies, a total of 5,287 genes were differentially expressed between control and stress conditions (Figure 5). This number was much lower for NC colonies with 1,460 differentially expressed genes (adjusted *P* < 0.05).

**Figure 5.**
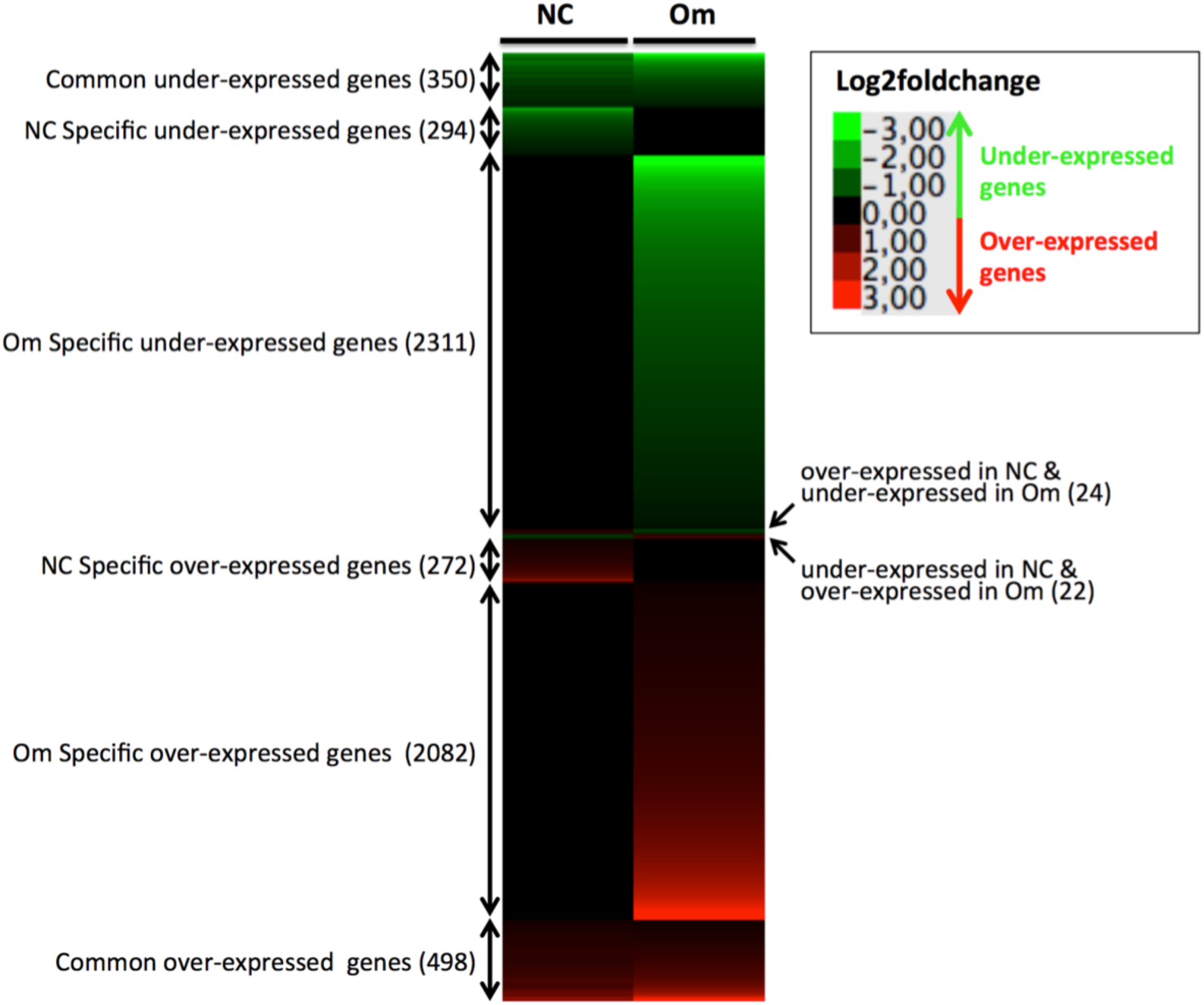
Heatmap and clustering of significantly differentially expressed genes between the control and the heat stress condition for colonies from each locality. Each gene is represented by a row.

Among differentially expressed genes, 848 were differentially expressed in the same direction in both localities (498 over-expressed and 350 under-expressed). In both cases, the differential expression level was significantly higher for the Om corals with a mean log2-fold change of 0.9 for shared over-expressed genes in Om *vs*.

0.6 in NC (Wilcoxon test; *P* < 0.0001), and −1.2 for the shared under-expressed genes in Om *vs*. −0.8 in NC (Wilcoxon test; *P* < 0.0001) (Figure 6 and Supplementary Table S8).

**Figure 6.**
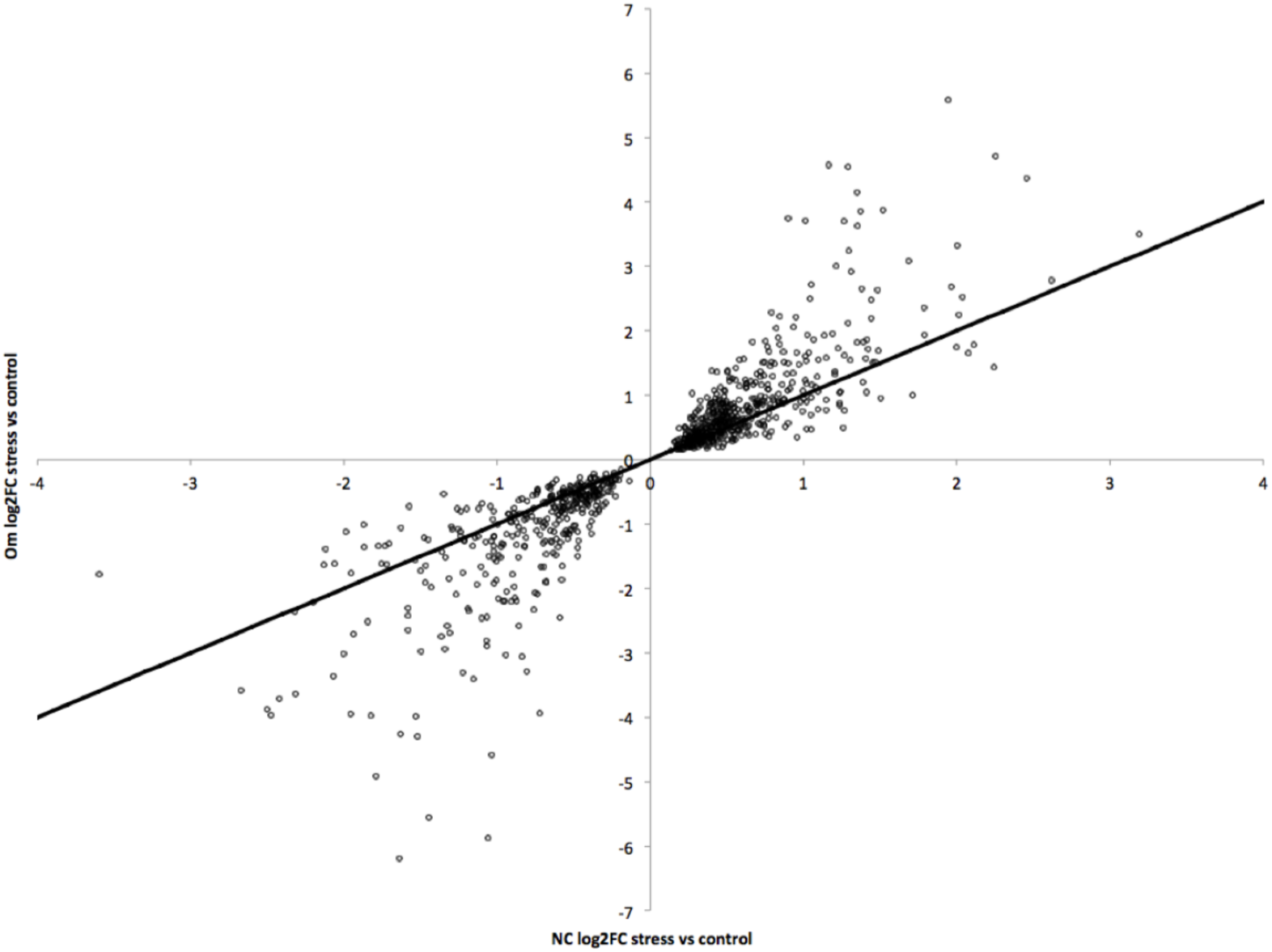
Scatterplot of the log2-fold changes in gene expression in response to heat stress in the Om colonies (y-axis) vs. the NC colonies (x-axis) for the 848 genes that were over-expressed (498 genes) or under-expressed (350 genes) in colonies from both localities. The line represents the y=x line depicting similar responses between colonies.

Additionally, colonies from the two localities also responded specifically to heat stress. In particular, 272 genes were over-expressed and 294 were under-expressed only in the NC corals, whereas 2,082 were over-expressed and 2,311 were under-expressed only in the Om ones when exposed to heat stress. Finally, the colonies from both localities displayed antagonistic transcriptomic responses to heat stress for a small subset of genes (24 over-expressed in NC but under-expressed in Om, and 22 under-expressed in NC but over-expressed in Om).

Altogether these results revealed a greater transcriptomic response to heat stress in colonies originating from Oman compared to those from New Caledonia (4,393 differentially expressed genes for the Om corals *vs.* 566 genes for the NC ones).

### Discriminant Analysis of Principal Components (DAPC)

At the overall gene expression level, our DAPC analysis clearly discriminated the colonies from both localities (Figure 7). More interestingly, the GLM revealed a significant interaction term between the locality and condition (control or heat stress) effects (*P* = 0.04), hence indicating that the slope of the reaction norm was different between localities. More particularly, the Om colonies responded to a greater extent than the NC ones, and thus showed significantly higher gene expression plasticity in response to heat stress.

**Figure 7.**
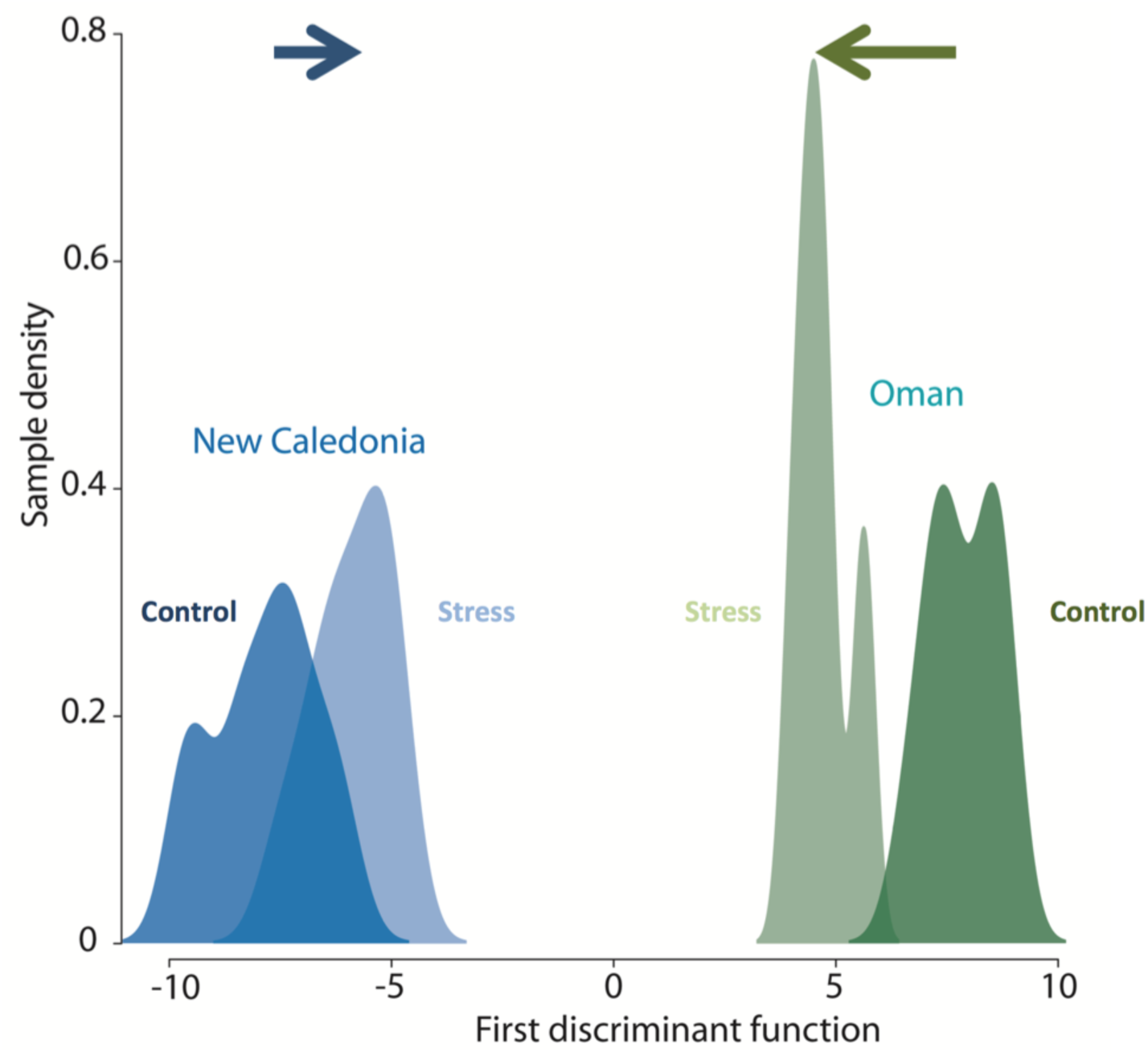
Colony level gene expression variation in response to heat stress, based on DAPC analysis. The x-axis is the first discriminant function of the DAPC along which the overall gene expression difference between colonies at both experimental conditions (stress and control) and from both localities (NC and Om) was maximized. This indicates the degree of similarity between the transcriptomes. The density plots obtained for NC and Om colonies are represented in blue and green, respectively. Dark and light density plots correspond to samples from the control and stress experimental conditions. The arrows above the density plots represent the direction of the mean change in the gene expression profiles.

It is worth stressing that colonies having experienced the heat stress displayed more similar genome-wide expression profiles than controlled colonies (Figure 7). Such pattern was also observed in colonies of mustard hill coral *P. astreoides* experiencing a heat stress compared to controls (Kenkel & Matz 2016). This apparent convergence in the functional response of colonies from different habitats to heat stress might be at least partly explain by the fact some common molecular pathways are turned-on when colonies are facing stressful conditions although the magnitude of such responses is different.

### Analysis of gene function

To investigate the functions associated with the differentially expressed genes we performed a BLASTX annotation of transcripts followed by a Gene Ontology (GO) term (biological process, molecular function, and cell compartment) (Supplementary File S9).

For the 498 common over-expressed genes, 139 biological processes were enriched compared to the entire set of annotated genes. The most significant biological process identified in the REVIGO analysis (i.e. with lowest FDR= 2.1×10^-68^) was response to stress (Figure 8). Following this sequentially, were cellular metabolism (FDR=3.7×10^-49^), positive regulation of biological processes (FDR = 2.4×10^-43^), cell death (FDR = 2.5×10^-33^), cellular localization (FDR = 8.4×10^-25^), and pigment metabolism (FDR = 2.1×10^-21^). Among the 272 genes over-expressed in the NC but not in the Om colonies in response to heat stress, 38 biological processes were enriched: organic acid catabolism (FDR = 1.6×10^-22^), protein transport (FDR = 1.8×10^-16^), stress response (FDR = 4.8×10^-13^), and cellular metabolism (FDR = 3×10^-12^) were the four most significantly enriched biological processes (Figure 8). Among the 2,082 genes over-expressed in the Om but not in the NC colonies in response to heat stress, 160 biological processes were enriched, the most significant being ncRNA metabolism (FDR = 8.9×10^-303^), cellular metabolism (FDR = 4.4×10^-70^), carbohydrate derivative biosynthesis (FDR = 5.9×10^-64^), and organic substance transport (FDR = 2×10^-44^).

**Figure 8.**
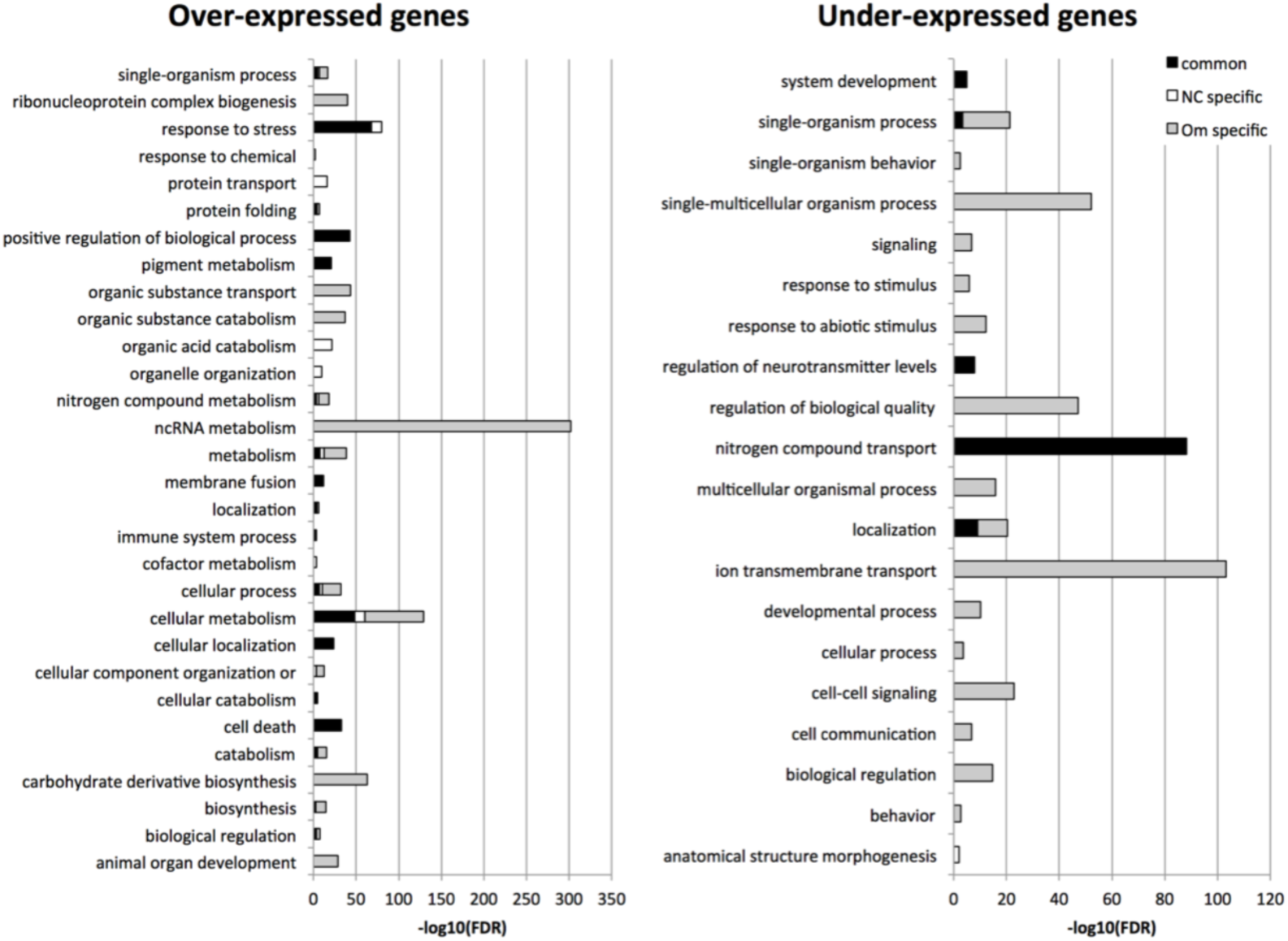
Summary of the GO enrichment analysis following REVIGO synthesis. Each enriched biological process is represented by a bar proportional to the log10(FDR). The colors correspond to the three categories of genes (common: black; Om-specific: grey; NC-specific: white) that were over-expressed (left panel) or under-expressed (right panel).

For the 350 genes that were under-expressed following heat stress irrespective to the locality of origin (Om or NC), 48 biological processes were enriched and grouped into five biological processes: nitrogen compound transport (FDR = 5.4×10^-89^), localization (FDR = 8.1×10^-10^), regulation of neurotransmitter levels (FDR = 1.2×10^-8^), system development (FDR = 8.8×10^-6^), and single organism process (FDR = 4×10^-4^). Among the under-expressed genes in the NC colonies only, a single biological process (anatomical structure/morphogenesis) was found to be enriched (FDR = 9×10^-3^). Among the under-expressed genes in the Om colonies, 139 biological processes were enriched, with the most significant being ion transmembrane transport (FDR = 7.6×10^-104^), single multicellular organism process (FDR = 7.5×10^-53^), regulation of biological quality (FDR = 6×10^-48^), cell–cell signaling (FDR = 1.5×10^-23^), single organism process (FDR = 1.1×10^-18^), multicellular organism process (FDR = 1.5×10^-16^), biological regulation (FDR = 2.3×10^-15^), response to abiotic stimulus (FDR = 6.2×10^-13^), and localization (FDR = 4.6×10^-12^). The complete results for all GO term categories including molecular function are available in Supplementary File S9.

Regarding cellular compartments, the mitochondria was the most significantly enriched in the common over-expressed genes (FDR = 1.5×10^-180^), as well as among genes over-expressed in NC (FDR = 2.5×10^-82^) while in genes over-expressed in Om corals the intracellular organelle lumen was the most significantly enriched (FDR = 1×10^-560^).

To investigate whether the presumably more thermotolerant colonies from Oman displayed a frontloading strategy (i.e. a higher expression for some genes compared to the colonies from NC) as previously described in scleractinian corals (Barshis *et al*. 2013), we compared the gene expression levels in control conditions between Om and NC colonies for those genes that were over-expressed in NC colonies (Supplementary File S10). This comparison revealed that the constitutive expression level was often greater in the Om colonies. Among the 770 genes that were over-expressed in NC colonies in response to thermal stress (272 specifically and 498 in common with Om), 484 were constitutively (i.e. in the control condition) more expressed in Om. Among these genes, 163 were not differentially expressed between the control and stress temperatures reflecting true frontloading based on the definition of Barshis et al. (2013), while 301 were over-expressed and only 20 were under-expressed during heat stress. These 484 genes with higher constitutive expression in Om were submitted to GO term enrichment analysis. No significant results were found for the under-expressed genes. The frontloaded genes were enriched in the biological processes cellular respiration (FDR = 4.4×10^-23^), cellular component organization (FDR = 0.002), homeostatic process (FDR = 0.005), cellular component organization or biogenesis (FDR = 0.007), cofactor metabolism (FDR = 0.009), and stress response (FDR = 0.009), and in the mitochondrion for the most significant cellular compartment (FDR = 1.6×10^-66^). Most interestingly, for genes associated with a higher basal expression level together with over-expression in the Om colonies, the most enriched biological processes were stress response (FDR = 1.2×10^-26^), pigment metabolism (FDR = 5.1×10^-24^), regulation of phosphate metabolism (FDR = 3.2×10^-15^), cellular metabolism (FDR = 2.7×10^-11^), and protein folding (FDR = 7.3×10^-6^).

## Discussion

### Specific context of adaptation

Our aim was to compare the phenotypic plasticity in terms of transcriptomic response to heat stress of coral colonies originating from different localities displaying contrasted thermal regimes. As morphology can be misleading for species identification in scleractinians, notably in *Pocillopora* genus (Gélin et al., 2017a), we used a molecular approach to test the species relationships of our samples. The analysis of mitochondrial sequences and clustering analyses indicated that, despite similar morphologies, our samples corresponded to different species. This agrees well with previous works showing the importance of cryptic lineages and morphological plasticity in the *Pocillopora* genus (Gélin *et al*. 2017a and references herein). Oman colonies corresponded to species hypothesis PSH12 of (Gélin *et al*. 2017b), which is restricted to the Northwestern Indian Ocean. Regarding the two species hypotheses from NC, SSH05a (*P. damicornis* type *β* SSH05a or *P. acuta*) is found in the Pacific Ocean and PSH04 (*P. damicornis* type *α* or *P. damicornis sensu stricto*) is nearly exclusively found in the Pacific Ocean (very rare in the Indian Ocean, and not found yet in Red Sea) (Gélin *et al*. 2017b). It would be interesting to study whether inside each species hypothesis, different thermotolerance phenotypes are present. Conversely, the observation of a similar response to thermal stress in two different species in NC, as revealed by differential gene expression as well as DAPC analyses, could indicate either a conserved strategy or a convergence under the same ecological conditions.

### An ecologically realistic heat stress

The heat stress applied in this study was ecologically realistic, since the first visual response (i.e. polyp closure) was observed for all colonies when the gradually increasing experimental temperature reached the upper temperature they are subjected to *in natura* (30°C and 34°C for NC and Om corals, respectively). From a biological point of view this first result hence clearly supports that these colonies from two localities that are experiencing two different thermal regimes *in natura* display differential ability to deal with heat stress. Moreover, the accurate control of all other seawater parameters allows us to consider that the holobiont response to the thermal treatment is specific to heat stress and not to other possible confounding effects. Last, as we analyzed the samples before the first visible signs of stress (polyp closure), any change in the holobiont would therefore reflect the response to the heat stress and not homeostasis breakdown after disruption of the coral integrity.

### Symbiotic community: bacterial and Symbionaceae composition

For the bacterial community, we identified significant differences between localities and colonies. The microbiota composition of all samples was consistent with previous studies, showing a high proportion of Gammaproteobacteria and dominance of the symbiotic *Endozoicomonas* genus (Bourne & Munn 2005; Neave *et al*. 2016a; Peixoto *et al*. 2017). However, our results clearly demonstrate that neither maintenance in the experimental structure nor experimental heat stress induced major bacterial community changes in coral colonies irrespective to their locality of origin. For the Symbionaceae community, the ITS2 metabarcoding analysis enabled inter-clade resolution (Quigley *et al*. 2014). Two distinct types of D1a and C1 clades dominated, representing most of the sequences in the Om and NC corals, respectively. Nine ITS types (A to I) have been identified in the former genus *Symbiodinium* (Baker 2003). Some Symbiodiniaceae strains strongly participate to the overall holobiont fitness, with type D providing tolerance to higher temperatures (Berkelmans & van Oppen 2006) and C1 enhancing coral growth rates (Little *et al*. 2004). Interestingly, we found that the type D1a is dominant in the more thermotolerant Om corals, which is consistent with the results of previous works (Berkelmans & van Oppen 2006), however recent results shows that such an association is rather linked with minimal temperatures than annual amplitude of temperature changes (Brener-Raffalli *et al*. 2018).

Although the microbial community (both bacterial and Symbiodiniaceae) differed between the NC and Om corals, the composition did not change during transition from the field to the artificial seawater conditions, and remained similar during the experimental temperature increase. Thus, the coral holobiont assemblage remained similar after the course of the experiment. Such stability of the microbial community during experimental heat stress was previously observed in the scleractinian *Acropora millepora* (Bellantuono *et al*. 2012b) and *A. tenuis* (Littman *et al*. 2010). Thus, our study conforms to the idea that microbial communities associated with scleractinian corals remain unchanged when the holobionts are exposed to stressful temperatures (but see (Ziegler *et al*. 2017)) but further analyses of gene expression level would be needed to assess their functional responses. RNA-sequencing of eukaryotic poly-adenylated mRNA would allow in principle dual analysis of Symbiodiniaceae and coral host transcripts (Mayfield *et al*. 2014), but since our RNA extraction method resulted in very few algal transcripts, we only focused on the host transcriptomic response.

Based on these results, we investigated changes in host gene expression as the main mechanism of response to heat stress in our experimental design.

### Host transcriptomic response

Given the observed stability of the microbial symbiotic community during heat stress, we focused more specifically on the responses attributable to the coral host. We thus compared gene expression patterns at the qualitative and quantitative levels in Om and NC colonies in response to heat stress compared to the control condition. Altogether, our results clearly highlight that the Oman colonies exposed to more variable thermal conditions *in natura* also display, in response to heat stress, a greater plasticity in gene expression levels than the NC colonies. In particular, the transcriptomic response of the Oman colonies involved a larger number of genes with 73% of commonly differentially expressed genes having higher fold changes compared to the NC colonies. These findings are consistent with the theoretical expectations that a more variable environment promotes the evolution of a greater plasticity (Lande 2009). Accordingly, a recent transplantation study conducted *in natura* also identified greater transcriptomic plasticity in a more thermotolerant (in-shore) population compared with an (off-shore) population inhabiting a more stable thermal habitat in the mustard hill coral *P. astreoides* (Kenkel & Matz 2016).

Importantly however, we also identified several genes whose expression is constitutively higher in the Om colonies compared to the NC colonies by comparing the expression levels in the control condition. This process recently called “frontloading” (Barshis *et al*. 2013) reflects the preemptive expression of stress-response genes, hence predisposing organisms to better respond to stress. It has been proposed that the occurrence of plasticity *vs*. frontloading strategies may depend on the frequency of stresses relative to the typical response time of organisms, with frequent stresses promoting frontloading strategies whereas less frequent perturbations would result in an increased plasticity (Kenkel & Matz 2016). Other conceptual considerations especially in regards to the predictability of environmental variation through generations should also be taken into account (Danchin 2013; Herman *et al*. 2014). The frontloading is by definition more costly than plasticity since it transforms a response to the environment in a constitutive function. Frontloading is therefore a strategy that would be more efficient when offspring’s habitat is highly predictable. On the contrary, an unpredictable or less predictable offspring environment may promote plasticity to enable the exploration of a wider phenotypic landscape at a lesser cost. Plasticity and frontloading are often discussed as mutually exclusive responses (Barshis *et al*. 2013; Kenkel & Matz 2016). However, corals are known to display a high level of variation in their reproduction strategies (brooder vs. broadcast spawner) (Whitaker 2006; Baird *et al*. 2009), timing (Fan *et al*. 2006) and pelagic larval duration (Harrison & Wallace 1990). Environmental predictability in terms of stress frequency and annual temperature variation should be therefore limited and we hypothesized that, rather than being exclusive, plasticity and frontloading often co-occur especially in the population experiencing extreme environments.

Our results clearly support that plasticity and frontloading indeed co-occur specifically in the thermotolerant Om colonies experiencing a more variable thermal environment *in natura*. To tease apart the biological processes that are regulated via plasticity or frontloading in *Pocillopora* response to heat stress, we conducted an enrichment analysis. Keeping in mind that congruency between gene expression and protein levels should be cautious (Mayfield *et al*. 2016), we propose a detailed discussion of the response of coral colonies at the molecular level for each main biological process identified (Supplementary File S11). Notably, we found differences in gene expression levels in response to temperature increase between the two localities for genes involved in response to heat stress (such as HSPs), detoxification of reactive oxygen species, apoptosis, mitochondria energetic functioning, and symbiont maintenance with higher number of differentially expressed genes for the Om corals associated to higher fold changes. In corals, Hsp70 which displayed more intense overexpression in Om colonies, is one of the most documented protein chaperones associated to heat stress response (Barshis *et al*. 2013; Haguenauer *et al*. 2013). Several genes involved in oxidative stress had greater differential expression levels in Om such as calmodulin, quinone oxidoreductases and thioredoxin (Supplementary File S11). Reactive oxygen species (ROS), generated in consequence to heat stress, play a central role in host-symbiont interactions ultimately leading to bleaching as a host innate immune response to a compromised symbiont (see (Weis 2008) for a review). We also identified many genes involved in apoptosis, a process that has been recurrently associated with coral responses to heat stress (Ainsworth *et al*. 2011; Barshis *et al*. 2013; Pratlong *et al*. 2015; Maor-Landaw & Levy 2016).

Our results also suggest that allocating energy in heat stress response is at the expense of other crucial biological processes such as growth and reproductive functions as already shown in *Pocillopora* (Vidal-Dupiol *et al*. 2009; 2014), even if we could not test experimentally fitness effect of the experimental heat stress. However, the molecular mechanisms underlying such overall response to heat stress are still partly unresolved. Interestingly, we also found specific gene expression patterns linked with epigenetic regulation such as histone modifying enzymes as well as enzymes involved in DNA methylation and dsRNA regulation (Supplementary File S11). We also identified expression of many retro-transposons in Om. These processes could be involved in rapid epigenome modifications and thus fuel rapid adaptive evolution (Maumus *et al*. 2009; Torda *et al*. 2017; Jablonka 2017).

### Conclusion

Comparison of the response to an ecologically realistic heat stress of corals from the same genus but pertaining to different species hypotheses thriving in two contrasting thermal environments sheds light on the molecular basis of thermotolerance. We found that after heat exposure, the symbiotic community composition remained similar in colonies from both localities, but we identified major differences in gene regulation processes in the coral, thereby underlining the role of the coral host in the response to heat stress. The colonies from the locality displaying the most variable environment displayed (i) a more plastic transcriptome response involving more differentially expressed genes and higher fold expression changes; as well as (ii) a constitutive higher level of expression for another set of genes (frontloading). In the context of climate change, which is predicted to cause abnormal and rapid temperature increase (IPCC 2014), phenotypic plasticity and the capacity for rapid adaptation through epigenetic regulation and/or genetic assimilation would increase the probability of coral survival. Previous studies highlighted the importance of reef managements measures (Rogers *et al*. 2015) and assisted evolution (van Oppen *et al*. 2015), but also underlined the importance of preserving standing genetic/epigenetic variation in wild coral populations (Matz *et al*. 2017). Although the molecular mechanisms we described are most likely largely shared in this group of scleractinians, the question remains of the determinism of this thermotolerant phenotype and of the heritability of this character. It is however essential to keep in mind that even the most thermotolerant corals may bleach if they are exposed to temperature significantly higher to their own norm (Hughes *et al*. 2017b; Le Nohaïc *et al*. 2017).

## Data accessibility

The datasets generated and analyzed during the current study have been submitted to the SRA repository under bioproject number PRJNA399069 (https://www.ncbi.nlm.nih.gov/bioproject/PRJNA399069).

## Supplementary material

R Scripts are available online: https://osf.io/s5y34

## Acknowledgements

We are thankful to Dr. Madjid Delghandi from the Center of Marine Biotechnology at Sultan Qaboos University, the Al-Hail field station and the central laboratory of the College of Agricultural and Marine Sciences for providing us with necessary equipment during field work in Oman. We acknowledge Dr. Gilles Le Moullac and Dr. Yannick Gueguen from the Centre Ifremer du Pacifique for providing us aquaculture facilities during sampling. We acknowledge the Plateforme Gentyane of the Institut National de la Recherche Agronomique (INRA, Clermont-Ferrand, France) for microsatellite genotyping. We also thank the Genotoul bioinformatics platform, and the Toulouse Midi-Pyrenees and Sigenae group for providing help and computing resources (Galaxy instance; http://sigenae-workbench.toulouse.inra.fr as well as the Bio-Environnement platform (University of Perpignan) for sequencing and bioinformatics service. This project was funded by the ADACNI program of the French national research agency (ANR) (project no. ANR-12-ADAP-0016; http://adacni.imbe.fr), the Campus France PHC program Maïmonide-Israel and the DHOF program of the UMR5244 IHPE (http://ihpe.univ-perp.fr/en/ihpe-transversal-holobiont/). This work is a contribution to the Labex OT-Med (n° ANR-11-LABX-0061) funded by the ANR “Investissements d’Avenir” program through the A*MIDEX project (n° ANR-11-IDEX-0001-02). This study is set within the framework of the “Laboratoire d’Excellence (LABEX)” TULIP (ANR-10-LABX-41). We would like to thank Staffan Jacob, Xavier Pichon and Mar Sobral for their constructive and helpful comments. Version 4 of this preprint has been reviewed and recommended by Peer Community In Ecology (https://doi.org/10.24072/pci.ecology.100028).

## Appendix

**Supplementary Figure S1.** Experimental setup. Four tanks were used for each locality, 3 tanks containing the sampled colonies (one replicate per timepoint and per tank) and one additional tank as a control of coral health at the control temperature during the experiment.

**Supplementary Table S2.** Haplotype analysis of the six sampled colonies with microsatellite genotyping for the colonies from New Caledonia.

**Supplementary Figure S3.** Bacterial class composition (for the 24 most abundant) within each replicate for the Om and NC colonies, the three colonies of each locality, and three experimental conditions per colony. In situ (dark arrows); control temperature (green arrows); stress temperature (red arrows).

**Supplementary Table S4.** MANOVA results for beta diversity (Bray-Curtis distance) between localities, colonies, or experimental conditions.

**Supplementary File S5.** List and sequences of the 26,600 genes (XLOC) generated during RNAseq alignment.

**Supplementary Figure S6.** Heatmap and clustering of significantly differentially expressed genes between the control and the heat stress condition for each colony from the two localities. Each gene is represented by a line.

**Supplementary File S7.** DESeq2 results for the log2-fold changes, and adjusted p values between stress and control conditions for each locality (sheet 1) and for each colony (sheet 2).

**Supplementary Table S8.** Comparison between the log2-foldchange in Om and NC colonies for genes differentially under-expressed or over-expressed in the same way in colonies from both localities.

**Supplementary File S9.** GO enrichment results for biological processes, molecular functions, and cellular compartments for common, New Caledonia-specific, or Oman-specific over-expressed and under-expressed genes.

**Supplementary File S10.** Frontloaded genes in Oman corals among genes over-expressed in New Caledonia corals.

**Supplementary File S11.** Description of the functional analysis of genes, biological functions and cell compartment involved in the response to stress.

