## Supplementary figures and images for "Gene expression plasticity and frontloading promote thermotolerance in *Pocillopora* corals"

### Supplementary Figure S1

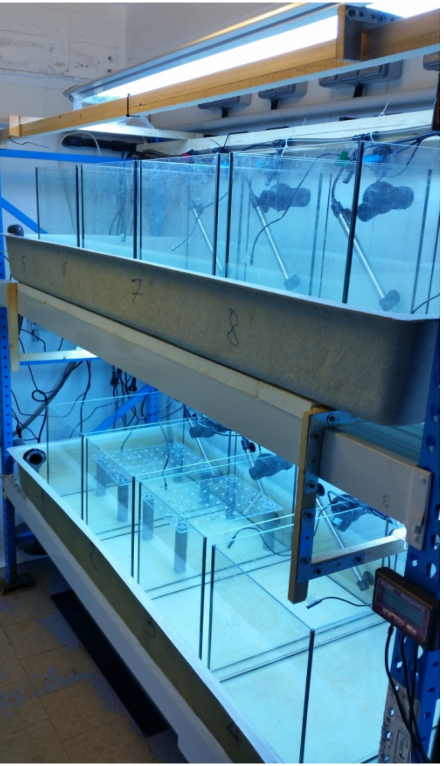

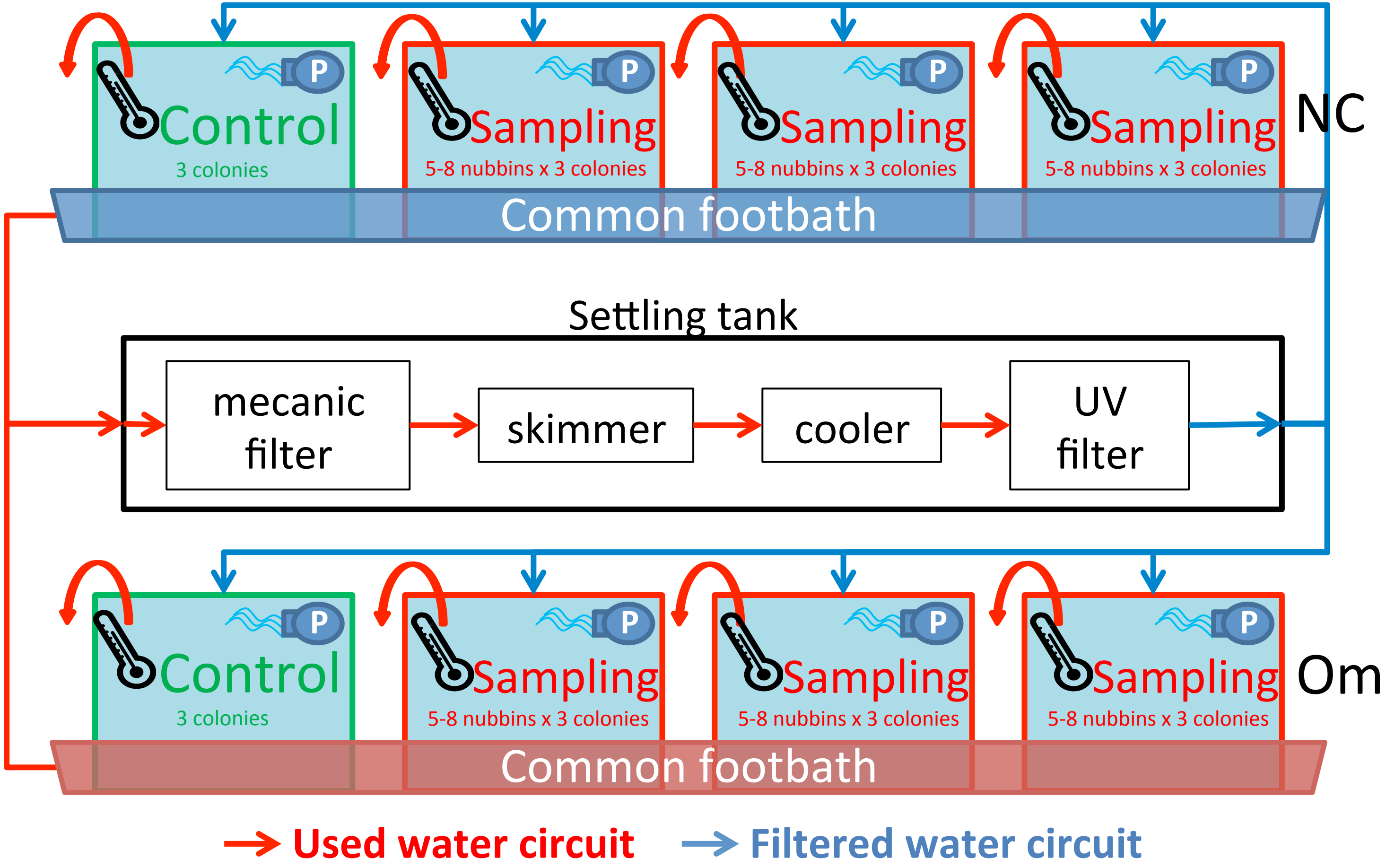

### Supplementary Figure S3

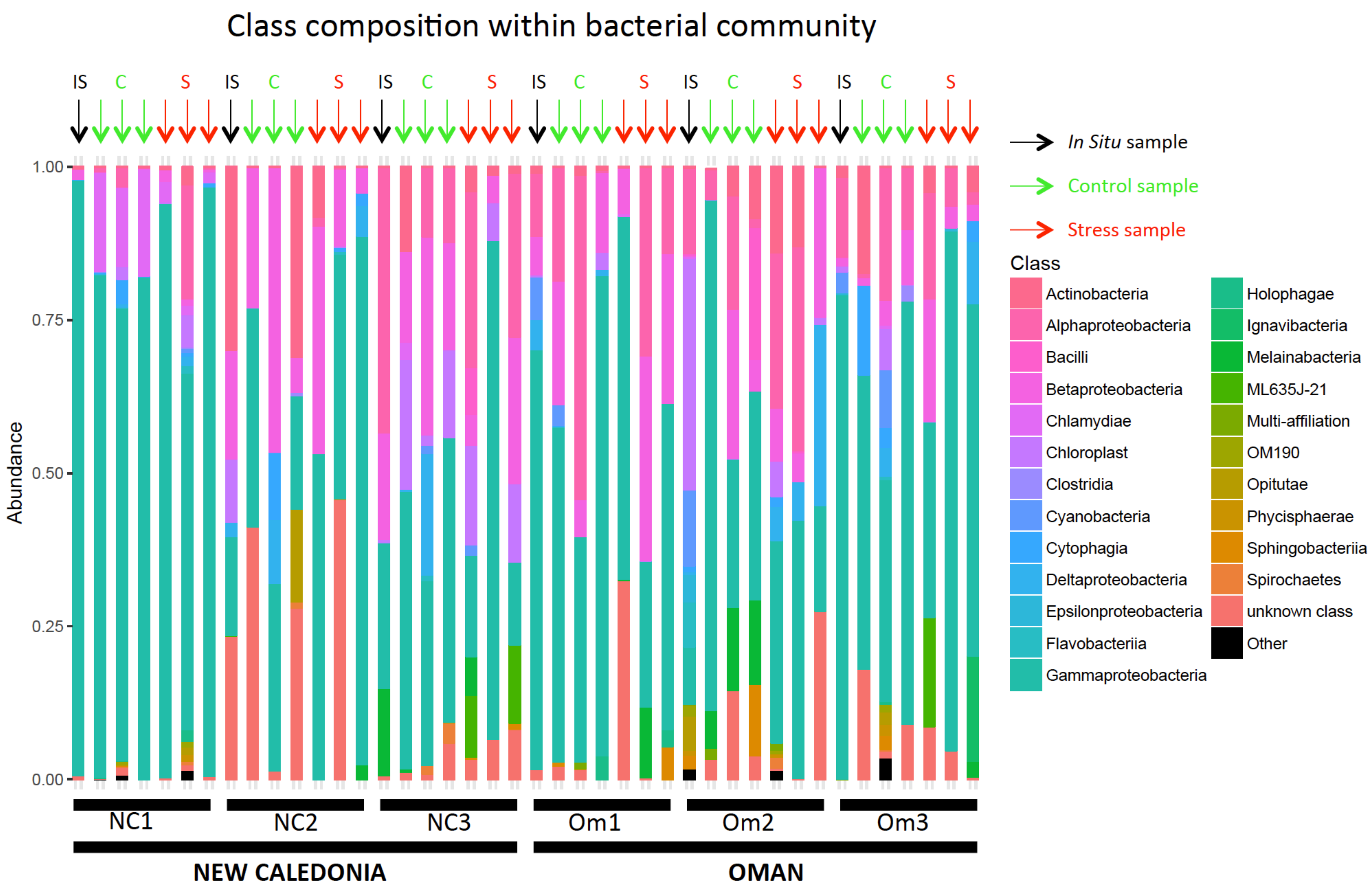

### Supplementary Figure S6

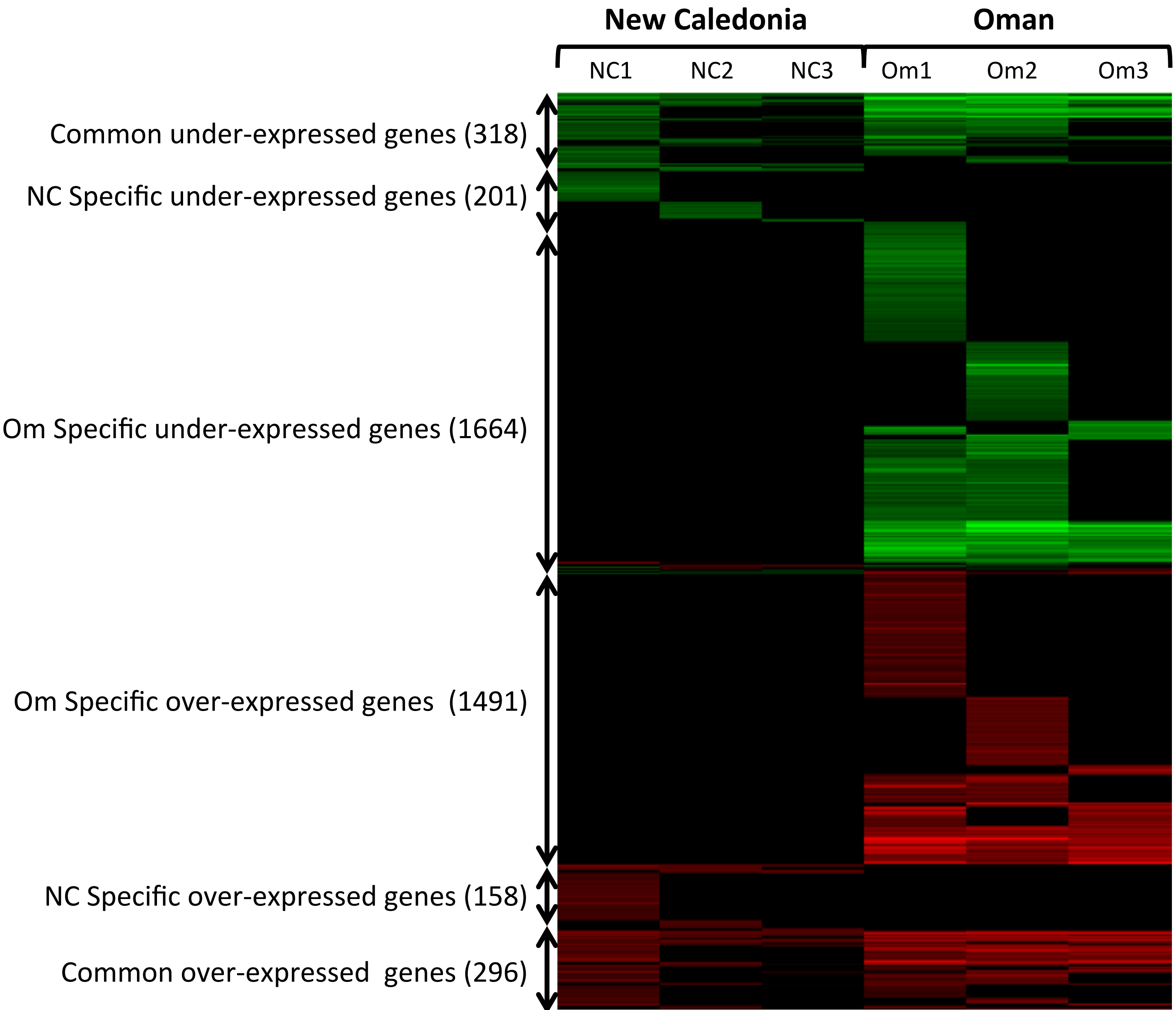
