## Supplementary File S11 for "Gene expression plasticity and frontloading promote thermotolerance in *Pocillopora* corals"

### Functional analyses

###### Response to stress:

###### HSPs:

Heat shock proteins (HSPs) are ubiquitous stress-induced chaperone molecules involved in protein folding and protein damage repair in most organisms (Feder & Hofmann, 1999). As such, several HSPs were found to be over-expressed in colonies from both localities, whichever their thermal regime, in response to heat stress. Among these, the over-expression of Hsp70 genes was significantly more intense in the Om colonies compared to NC ones. Hsp70 is one of the most documented protein chaperones in coral heat stress response (Barshis et al., 2013) (Haguenauer, Zuberer, Ledoux, & Aurelle, 2013), but other forms of HSPs such as Hsp60 (Brown, Downs, Dunne, & Gibb, 2002), Hsp90 (Carpenter, Patterson, & Bromage, 2010), and Hsp40 (Maor-Landaw & Levy, 2016) are also known to be involved in the response of these organisms. Accordingly we found that Hsp105, Hsp75, Hsp90, and Hsp71 were over-expressed only in Om corals and Hsp60 and Hsp71 showed higher basal expression levels in the Om corals. We also found that chaperones of the Hsp40 family (also known as DNAj) were regulated in response to heat stress; three of these were over-expressed in colonies from both localities with a higher basal expression level in the Om corals, whereas five were over-expressed only in the Om corals. These proteins seem to be implicated in protein folding during early heat stress in the scleractinian *Stylophora pistillata*, and are highly expressed prior to bleaching and tissue peeling (Maor-Landaw et al., 2014). We also found specific Om over-expression of the pyrexia gene, which codes for a transient receptor-activated cation channel, and had been shown to protect flies from high temperatures (Lee et al., 2005). Together these results indicate that most genes involved in the heat shock protein pathways are plastically regulated by corals when exposed to stress with a greater response in colonies inhabiting thermally variable environment. However few HSP genes are also frontloaded in these pre-exposed colonies, in particular Hsp60 which gene is encoded in the mitochondria. This pattern is likely to reflect a more general higher mitochondrial activity in colonies accustomed to more variable and/or warmer temperatures.

###### Reactive Oxygen Species (ROS) detoxification and dna repair:

One of the first consequences of heat stress is the production of reactive oxygen species (ROS), which might cause protein, membrane and DNA damages (Kowaltowski & Vercesi, 1999) (Yakovleva et al., 2009) (Jena, 2012). These oxidative molecules are produced in excess by the symbionts and mitochondria during the photosynthesis and the respiration processes, respectively. Cells produce several detoxification enzymes to limit cellular damages (Weis, 2008). In particular, under-expression of the calcium-binding messenger protein calmodulin is a sign of response to oxidative stress (Schallreuter, Gibbons, Zothner, Abou Elloof, & Wood, 2007) (Voolstra et al., 2009). This protein had more isoforms that were differentially expressed in response to heat stress in Om compared to NC corals, together with one calcium calmodulin-dependent kinase. The Quinone oxidoreductases proteins are involved in the generation of ROS (Porté et al., 2009) and were also specifically under-expressed in Om corals in response to heat stress. Conversely, the thioredoxin gene was over-expressed in colonies from both localities, but with a higher fold change in Om corals. This enzyme detoxifies oxidized molecules and is often implicated in coral response to heat stress (DeSalvo et al., 2010) (Maor-Landaw et al., 2014). Finally, polyamine oxidase is a key component of the oxidative burst in plants, and is involved in the induction of apoptosis (Yoda, Hiroi, & Sano, 2006). Polyamine oxidase was differentially expressed in response to heat stress in our experiment, under-expressed in Om corals and over-expressed in NC corals. Overall, the molecular pathway underlying ROS detoxification hence appears to be plastically regulated, with greater plasticity in the thermo-tolerant Om colonies than in the thermo-sensitive NC colonies.

Several genes coding for DNA repair enzymes including DNA excision repair, DNA ligase, and DNA polymerase were over-expressed in colonies from both localities when exposed to heat stress. However, among these genes, three showed higher basal expression levels and two were frontloaded in the Om colonies.

###### Apoptosis:

Among the genes differentially regulated in colonies when exposed to heat stress, many are also involved in apoptosis, a process that has been recurrently associated with coral responses to heat stress (Ainsworth et al., 2011) (Pratlong et al., 2015). Among these, TNFR and TRAF genes code for receptors and receptor-associated components of the tumor necrosis factor (TNF), the latter being central in the apoptosis pathway (Ashkenazi & Dixit, 1998). Five TNFR and 11 TRAF genes were specifically over-expressed in the Om stressed colonies, indicating a more intense regulation of the apoptosis pathway in these thermotolerant colonies. Furthermore, four TNFs were found to be specifically under-expressed in the Om corals, whereas TNFAIP3 was specifically over-expressed in response to heat stress. It is worth stressing that the TNFAIP3 gene (TNF alpha-induced protein3) is known to inhibit the NFkB process (inflammatory response) together with apoptosis (Opipari, Boguski, & Dixit, 1990). Moreover, Caspase8 and Caspase3 were specifically over-expressed in the Om stressed corals. Caspases constitute the effector core of the apoptotic process (Nicholson & Thornberry, 1997), and Caspase3 has commonly been implicated in the heat stress response in corals (Maor-Landaw & Levy, 2016). This further suggests that the thermotolerant Om colonies are capable of greater regulation of apoptosis than the thermosensitive NC ones when exposed to warm temperature. Another set of 10 TNFR genes and 3 TRAF genes were found to be over-expressed in colonies from both localities, but with a higher expression level in the Om corals both in the control and the stress conditions for 10 of the 13 genes irrespective to the treatment (i.e. stress and control temperature). Fem-1 homolog B, which is involved in the apoptotic process as a death receptor-associated protein, was also constitutively higher expressed in the Om compared to the NC colonies. Together these results indicate that part of the apoptosis pathway is frontloaded in the thermotolerant Om colonies. These results echoes with previous studies showing that TNFR genes are frontloaded in the common reef-building coral *Acropora hyacinthus* (Barshis et al., 2013) and hence strengthen the idea that frontloading such biological pathway is an adaptive response to variable and/or warmer temperatures in corals.

###### Energetic metabolism:

###### Mitochondrial functions:

The mitochondria are central to many important biological processes of their cell hosts including energy-generating respiration and can be the source of several oxidative molecules involved in apoptosis. As such, heat stress is expected to promote severe changes in the expression levels of genes involved in the mitochondrial function in all living organisms (Guderley & St-Pierre, 2002). Accordingly, our enrichment analysis highlighted several mitochondrial genes (either mitochondria- or nuclear-encoded). Among those, some of them were specifically or more intensively regulated in Om corals, including genes coding for 2-oxoglutarate dehydrogenase (which catalyzes the conversion of 2-oxoglutarate to succinyl-CoA and CO_2_ in the Krebs cycle), and several monocarboxylate transporters for lactate or pyruvate, the latter being essential for the regulation of energy metabolism in the thermosensitive sea anemone *Aiptasia* (Halestrap & Meredith, 2004) (Lehnert et al., 2014). These results corroborate with the general higher gene expression plasticity observed in colonies from a population experiencing a more variable thermal regime (Om) compared to colonies from a population exposed to narrower thermal amplitudes (NC).

Nevertheless, several other genes that participate in energy metabolism displayed a frontloaded pattern in the Om colonies such as the succinate dehydrogenase [ubiquinone] flavo mitochondrial-like and the citrate synthase genes. Interestingly, the expression of the latter gene is often used as a reliable indication of organisms’ aerobic capacity and mitochondrial density (e.g. (Rivest & Hofmann, 2014) (Hawkins & Warner, 2017)). Thus, it is likely that the observed frontloading pattern of these mitochondrial genes (encoded in the mitochondrial genome) reflects a greater mitochondrial density in the Om compared to the NC colonies. According to the literature related to non-phototrophic symbiotic ectotherms such increase in mitochondrial density as an adaptive response to heat stress appears counterintuitive since the opposite pattern is generally observed

(Pörtner, 2002). However, elevated mitochondrial activity on cnidarian hosts have already been found to be associated with thermal-tolerance in corals and sea anemones (Dixon et al., 2015; Hawkins & Warner, 2017). Hawkins & Warner suggest that such specific pattern might be at least partly explained by a decrease in symbiotic algae density resulting from heat stress. Energy metabolism in scleractinian corals is highly dependent of their symbiotic algae, which are the major source of glucose generated by photosynthesis. A partial loss of their symbiotic algal may force cnidarian hosts to increase their energy metabolism, hence requiring an increase in their mitochondrial contents, and to use heterotrophic sources of energy that can be obtained from feeding or from the use of metabolic reserve such as lipids. In this context we also observed a high level of regulation of lipid metabolism in the Om corals. The apolipo A (a constituent of lipoprotein associated with lipid dissolution) and dLp HDL-BGBP precursor (which has lipid transporter activity) genes were specifically over-expressed in the Om corals, and could facilitate the transport and assimilation of lipids.

###### Symbiont regulation:

Perturbation of energy metabolism and a decrease in the efficiency of energy production by symbiotic algae can be linked to the modulation of genes associated with symbiont maintenance. In our experiment three Rab-11 isoforms were found to be over-expressed in colonies when exposed to stress, including: one being over-expressed in colonies from both localities but with a higher fold change in the Om corals; one that occurred in NC corals while showing frontloading expression in Om corals; and one induced only in the Om corals. This recycling regulatory protein has been shown to regulate phagosomes containing *Symbiodinium* cells in the *Aiptasia-Symbiodinium* model system (Chen et al., 2005). Calumenin was also over-expressed in both Om and NC colonies in response to heat stress, but had a higher fold change in Om corals. This protein is known to be a signal intermediate for sym32, a signaling protein involved in symbiont recognition by the coral host (Reynolds, Schwarz, & Weis, 2000; Schwarz & Weis, 2003). In addition, three lectins exhibited differential expression in the Om corals: one was over-expressed (with a log2-fold change of 4.1) and two were under-expressed (log2-fold change of -2.8 and -2.5). It is worth stressing that lectins are key proteins involved in symbiont recognition (Wood-Charlson, Hollingsworth, Krupp, & Weis, 2006) and that a breakdown of this recognition may lead to the disruption of the symbiosis (Vidal-Dupiol et al., 2009). Thus although we did not expressly quantify the density of *Symbiodinium* algae in colonies during our experiment, our results strongly suggest that *Symbiodinium* were at least partly excluded as a response to heat stress prior to bleaching process. These results hence corroborate with those found relative to a shift in both the greater involvement in energy metabolism and resources use shift.

###### Cnidocytes:

We observed a higher level of regulation of genes involved in the functioning of cnidocytes (also known as nematocytes) in Om corals. These ectodermal cells are specific to cnidarians, and are involved in environment sensing, defense, and predation. Three cytosolic phospholipase A2 (cPLA2) genes were specifically over-expressed in Om corals. In cnidocytes, the corresponding proteins are involved in venom efficacy by promoting prey lysis (Argiolas & Pisano, 1983) (Nevalainen et al., 2004). One MAC perforin domain-containing gene had a higher basal expression level in Om corals. This gene encodes for a membrane attack complex/perforin-domain containing protein, and is found in gland cells and nematocytes in the sea anemone *Nematostella*, suggesting a potential role in prey killing (Miller et al., 2007). Conversely, we found under-expression in Om corals of a gigantoxin homolog, which is a cytolysin (actinoporin) found abundantly in cnidocytes (Hu et al., 2011) (Frazão, Vasconcelos, & Antunes, 2012). Several polycystic kidney disease gene (PKD1 and 2) isoforms were also found to be under-expressed in the Om corals. These proteins are localized in tentacles in Hydra, and seem to be involved in cnidocyte discharge (McLaughlin, 2017). These changes could have been promoted by notch signaling (with over-expression of one notch homolog 2); in cnidarians, this pathway is particularly important for cnidogenesis as shown in *Nematostella vectensis* (Marlow, Roettinger, Boekhout, & Martindale, 2012). Together these results clearly indicate that the Om colonies are better adapted to switch to heterotrophic nutrition in response to heat stress. Such switch could be linked to higher levels of resistance to heat stress via a decrease in *Symbiodinium* cell numbers, as described in other scleractinian corals (Hughes & Grottoli, 2013) (Aichelman et al., 2016).

To summarize, we found that Om colonies displayed a more plastic transcriptomic response for many genes involved in fundamental biological pathways such as stress-response pathways (e.g. HSPs, apoptosis, DNA repair, ROS management) in response to heat stress. Moreover, several genes were found to be frontloaded especially in the apoptosis as well as in the energy metabolic pathway. The latter pattern could be explained by a higher regulation of metabolic genes but could also reflect in part an increase in mitochondrial activity as already documented in other symbiotic cnidarians in response to heat stress (Rivest & Hofmann, 2014). Higher plasticity and frontloading are two strategies that are costly. This suggests possible trade-offs with other critical life history traits such as growth and reproduction. In the next sections we will discuss the transcriptomic response of both Om and NC colonies in response to stress at molecular pathways associated with central fitness traits.

###### Morpho-anatomic processes and reproduction:

###### Skeleton and muscle cells:

Some enzymes potentially involved in the calcification and muscle contraction were under-expressed in response to heat stress in Om corals, in particular, a single chain carbonic anhydrase and four carbonic anhydrase genes (1, 2, 7, and 12). Carbonic anhydrases catalyze the interconversion of CO_2_ and water to bicarbonate and protons, enabling maintenance of the acid-base balance (Tashian, 1989). In scleractinian corals, carbonic anhydrases are also involved in biomineralization processes involved in skeleton formation (Bertucci et al., 2013). Two hephaestin genes were also under-expressed following heat stress in Om corals. These copper-dependent ferroxidases were identified in the skeleton of *Acropora millepora* (Vulpe et al., 1999), where they can be involved in the incorporation and regulation of iron in the carbonate skeleton (Ramos-Silva et al., 2013). Several myosin related proteins similar to myosin light chain kinase or myosin regulatory light polypeptide 9 were found to be specifically under-expressed in the Om corals. Actin-binding LIM partial, potentially involved in Z-disk activity of muscles, and several genes having the LIM domain were also found to be under-expressed in Om corals. In eumetazoans including cnidarians, these proteins are involved in regulation of muscle activity (for a review, see (Leclère & Röttinger, 2016). In addition, the genes for the acetyl choline receptor “43 kDa receptor-associated of the synapse-like” and the “calcium-activated potassium channel subunit beta-like” proteins, both of which down-regulate muscle contraction, were over-expressed in Om colonies, which is in accordance with a global decrease in functioning of the contractile apparatus. The observed down-regulation of skeleton formation and muscle contraction strongly suggest that mounting a higher –and most likely better– transcriptomic response to heat stress is associated with a decrease energy allocation to crucial fitness traits and particularly to growth in scleractinian corals.

###### Nervous system:

Many genes involved in nervous system functioning were differentially regulated, with some being over-expressed (e.g. neuronal acetylcholine receptor and VWFA, which regulate the calcium dependent voltage chain), and others being under-expressed, including a proton myo-inositol cotransporter, SCO-spondin (axon guidance), sodium- and chloride-dependent GABA and taurine transporters (neurotransmission), sodium-dependent phosphate transport 2B, and synaptotagmin. Many transcription factors that could be involved in morpho-anatomic integrity, including numerous forkhead box homologs implicated in cell growth, proliferation, and differentiation were over-expressed in Om corals. All these may be involved more globally in the regulation of morpho-anatomic processes (muscle contraction, cnidocytes).

CIRCADIAN CLOCK / REPRODUCTION:

Several photoreceptors were differentially expressed in the corals. A cryptochrome gene was frontloaded in Om colonies. These flavoproteins are found in plants, animals, insects, and cnidarians, and in corals it has been shown that cryptochrome can synchronize the circadian clock and the reproductive system (Reitzel, Tarrant, & Levy, 2013). Three melatonin receptors were also found to be regulated in response to heat stress in Om corals (two over-expressed and one under-expressed). As in vertebrates, in which melatonin is well characterized, in cnidarians it is also involved in the circadian clock (Peres et al., 2014). Similarly, two under-expressed and three over-expressed opsin-like genes were identified. The protein is a homolog of the photoreceptor melanopsin, and in *Acropora* is involved in circadian cycle regulation (Vize, 2009).

Taken together, changes in morpho-anatomic regulation and perturbations in circadian cycles and reproduction may also reflect a trade-off mechanism in the stress response of Om colonies, which is consistent with greater plasticity, and intensive and not fully specific gene regulation activity. We believe that our study contribute to a better comprehension of the biological processes involved in coral thermotolerance. In particular we highlighted the molecular pathways that were regulated either via frontloading or plasticity or both. Our results also suggest that allocating energy in heat stress response to better cope with higher temperatures is at the expense of other crucial biological processes such as growth and reproductive functions even if we could not test experimentally fitness effect of the ecologically realistic heat stress. However, the molecular mechanisms underlying such global response to heat stress are still partly unresolved. Interestingly we also found specific gene expression patterns linked with epigenetic regulation that could be involved in such mechanisms and which are discussed in the next and last section.

###### Epigenetic processes:

Two histone acetyltransferase genes were over-expressed in Om corals. These enzymes, which are also found in *Nematostella*, are involved in epigenetic gene regulation by modifying the nucleosome structure, and thus the transcription of genes (Sterner & Berger, 2000) (Karmodiya et al., 2014). Several histone–lysine N-methyltransferases were also identified, five of which were under-expressed (four PRDM6 and one setd3 isoform) and three of which were over-expressed in Om corals. As with histone acetyltransferase, these enzymes are involved in gene expression regulation through their influence on chromatin structure (Huang, 2002) (Vervoort, Meulemeester, Béhague, & Kerner, 2016). One lymphoid-specific helicase-like (HELLS) gene was frontloaded in Om corals. This protein is known to participate in epigenetic processes, and interacts with DNMT1 (cytosine methyl-transferase) for the maintenance of *de novo* methylation (Dabe, Sanford, Kohn, Bobkova, & Moroz, 2015; Myant et al., 2011). We identified under-expression of two SID-1 transmembrane family member genes involved in dsRNA regulation (W. Li, Koutmou, Leahy, & Li, 2015). Many reverse transcriptase homologues, including RNA-directed DNA polymerase from a jockey-like mobile element, were differentially expressed or more intensely regulated during heat stress in the Om corals. These reverse transcriptases are typical of the retro-transposon activity known to be activated or less controlled during general stress (Wessler, 1996). The loss of inhibition of retro-transposon activity may reflect impairment in the control as a result of possible trade-off with more specific functions, but could also be a “SOS-like” survival mechanism. Indeed, in a threatening environment a high transposition activity may induce mutations in polyps with potential genomic changes, which could fuel rapid adaptive evolution {Jablonka:2017kh, Maumus:2009gf}.

Aichelman, H. E., Townsend, J. E., Courtney, T. A., Baumann, J. H., Davies, S. W., & Castillo, K. D. (2016). Heterotrophy mitigates the response of the temperate coral Oculina arbuscula to temperature stress. *Ecology and Evolution*, *6*(18), 6758–6769. http://doi.org/10.1002/ece3.2399

Ainsworth, T. D., Wasmund, K., Ukani, L., Seneca, F., Yellowlees, D., Miller, D., & Leggat, W. (2011). Defining the tipping point: a complex cellular life/death balance in corals in response to stress. *Scientific Reports*, *1*, 160. http://doi.org/10.1038/srep00160

Argiolas, A., & Pisano, J. J. (1983). Facilitation of phospholipase A2 activity by mastoparans, a new class of mast cell degranulating peptides from wasp venom. *The Journal of Biological Chemistry*, *258*(22), 13697–13702.

Ashkenazi, A., & Dixit, V. M. (1998). Death receptors: signaling and modulation. *Science (New York, N.Y.)*, *281*(5381), 1305–1308.

Barshis, D. J., Barshis, D. J., Ladner, J. T., Ladner, J. T., Oliver, T. A., Oliver, T. A., et al. (2013). From the Cover: Genomic basis for coral resilience to climate change. *Proceedings of the National Academy of Sciences*, *110*(4), 1387–1392. http://doi.org/10.1073/pnas.1210224110

Bertucci, A., Moya, A., Tambutte, S., Allemand, D., Supuran, C. T., & Zoccola, D. (2013). Carbonic anhydrases in anthozoan corals-A review. *Bioorganic & Medicinal Chemistry*, *21*(6), 1437–1450. http://doi.org/10.1016/j.bmc.2012.10.024

Brown, B. E., Downs, C. A., Dunne, R. P., & Gibb, S. W. (2002). Exploring the basis of thermotolerance in the reef coral Goniastrea aspera. *Marine Ecology Progress Series*, *242*, 119–129. http://doi.org/10.3354/meps242119

Carpenter, L. W., Patterson, M. R., & Bromage, E. S. (2010). Water flow influences the spatiotemporal distribution of heat shock protein 70 within colonies of the scleractinian coral Montastrea annularis (Ellis and Solander, 1786) following heat stress: Implications for coral bleaching. *Journal of Experimental Marine Biology and Ecology*, *387*(1-2), 52–59. http://doi.org/10.1016/j.jembe.2010.02.019

Chen, M.-C., Hong, M.-C., Huang, Y.-S., Liu, M.-C., Cheng, Y.-M., & Fang, L.-S. (2005). ApRab11, a cnidarian homologue of the recycling regulatory protein Rab11, is involved in the establishment and maintenance of the Aiptasia-Symbiodinium endosymbiosis. *Biochemical and Biophysical Research Communications*, *338*(3), 1607–1616. http://doi.org/10.1016/j.bbrc.2005.10.133

Dabe, E. C., Sanford, R. S., Kohn, A. B., Bobkova, Y., & Moroz, L. L. (2015). DNA Methylation in Basal Metazoans: Insights from Ctenophores. *Integrative and Comparative Biology*, *55*(6), 1096–1110. http://doi.org/10.1093/icb/icv086

DeSalvo, M. K., Sunagawa, S., Fisher, P. L., Voolstra, C. R., Iglesias-Prieto, R., & Medina, M. (2010). Coral host transcriptomic states are correlated with Symbiodiniumgenotypes. *Molecular Ecology*, *19*(6), 1174–1186. http://doi.org/10.1111/j.1365-294X.2010.04534.x

Dixon, G. B., Davies, S. W., Aglyamova, G. A., Meyer, E., Bay, L. K., & Matz, M. V. (2015). CORAL REEFS. Genomic determinants of coral heat tolerance across latitudes. *Science (New York, N.Y.)*, *348*(6242), 1460–1462. http://doi.org/10.1126/science.1261224

Feder, M. E., & Hofmann, G. E. (1999). Heat-shock proteins, molecular chaperones, and the stress response: evolutionary and ecological physiology. *Annual Review of Physiology*, *61*, 243–282. http://doi.org/10.1146/annurev.physiol.61.1.243

Frazão, B., Vasconcelos, V., & Antunes, A. (2012). Sea anemone (Cnidaria, Anthozoa, Actiniaria) toxins: an overview. *Marine Drugs*, *10*(8), 1812–1851. http://doi.org/10.3390/md10081812

Guderley, H., & St-Pierre, J. (2002). Going with the flow or life in the fast lane: contrasting mitochondrial responses to thermal change. *The Journal of Experimental Biology*, *205*(Pt 15), 2237–2249.

Haguenauer, A., Zuberer, F., Ledoux, J.-B., & Aurelle, D. (2013). Adaptive abilities of the Mediterranean red coral Corallium rubrum in a heterogeneous and changing environment: from population to functional genetics. *Journal of Experimental Marine Biology and Ecology*, *449*, 349–357. http://doi.org/10.1016/j.jembe.2013.10.010

Halestrap, A. P., & Meredith, D. (2004). The SLC16 gene family-from monocarboxylate transporters (MCTs) to aromatic amino acid transporters and beyond. *Pflugers Archiv : European Journal of Physiology*, *447*(5), 619–628. http://doi.org/10.1007/s00424-003-1067-2

Hawkins, T. D., & Warner, M. E. (2017). Warm preconditioning protects against acute heat-induced respiratory dysfunction and delays bleaching in a symbiotic sea anemone. *The Journal of Experimental Biology*, *220*(6), 969–983. http://doi.org/10.1242/jeb.150391

Hu, B., Guo, W., Wang, L.-H., Wang, J.-G., Liu, X.-Y., & Jiao, B.-H. (2011). Purification and characterization of gigantoxin-4, a new actinoporin from the sea anemone Stichodactyla gigantea. *International Journal of Biological Sciences*, *7*(6), 729–739.

Huang, S. (2002). Histone methyltransferases, diet nutrients and tumour suppressors. *Nature Reviews Cancer*, *2*(6), 469–476. http://doi.org/10.1038/nrc819

Hughes, A. D., & Grottoli, A. G. (2013). Heterotrophic compensation: a possible mechanism for resilience of coral reefs to global warming or a sign of prolonged stress? *PloS One*, *8*(11), e81172. http://doi.org/10.1371/journal.pone.0081172

Jena, N. R. (2012). DNA damage by reactive species: Mechanisms, mutation and repair. *Journal of Biosciences*, *37*(3), 503–517.

Karmodiya, K., Anamika, K., Muley, V., Pradhan, S. J., Bhide, Y., & Galande, S. (2014). Camello, a novel family of Histone Acetyltransferases that acetylate histone H4 and is essential for zebrafish development. *Scientific Reports*, *4*, 6076. http://doi.org/10.1038/srep06076

Kowaltowski, A. J., & Vercesi, A. E. (1999). Mitochondrial damage induced by conditions of oxidative stress. *Free Radical Biology & Medicine*, *26*(3-4), 463–471.

Leclère, L., & Röttinger, E. (2016). Diversity of Cnidarian Muscles: Function, Anatomy, Development and Regeneration. *Frontiers in Cell and Developmental Biology*, *4*, 157. http://doi.org/10.3389/fcell.2016.00157

Lee, Y., Lee, Y., Lee, J., Bang, S., Hyun, S., Kang, J., et al. (2005). Pyrexia is a new thermal transient receptor potential channel endowing tolerance to high temperatures in Drosophila melanogaster. *Nature Genetics*, *37*(3), 305–310. http://doi.org/10.1038/ng1513

Lehnert, E. M., Mouchka, M. E., Burriesci, M. S., Gallo, N. D., Schwarz, J. A., & Pringle, J. R. (2014). Extensive differences in gene expression between symbiotic and aposymbiotic cnidarians. *G3&#58; Genes|Genomes|Genetics*, *4*(2), 277–295. http://doi.org/10.1534/g3.113.009084

Li, W., Koutmou, K. S., Leahy, D. J., & Li, M. (2015). Systemic RNA Interference Deficiency-1 (SID-1) Extracellular Domain Selectively Binds Long Double-stranded RNA and Is Required for RNA Transport by SID-1. *The Journal of Biological Chemistry*, *290*(31), 18904–18913. http://doi.org/10.1074/jbc.M115.658864

Maor-Landaw, K., & Levy, O. (2016). Gene expression profiles during short-term heat stress; branching vs. massive Scleractinian corals of the Red Sea. *PeerJ*, *4*, e1814. http://doi.org/10.7717/peerj.1814

Maor-Landaw, K., Karako-Lampert, S., Waldman Ben-Asher, H., Goffredo, S., Falini, G., Dubinsky, Z., & Levy, O. (2014). Gene expression profiles during short-term heat stress in the red sea coral Stylophora pistillata. *Global Change Biology*, *20*(10), 3026–3035. http://doi.org/10.1111/gcb.12592

Marlow, H., Roettinger, E., Boekhout, M., & Martindale, M. Q. (2012). Functional roles of Notch signaling in the cnidarian Nematostella vectensis. *Developmental Biology*, *362*(2), 295–308. http://doi.org/10.1016/j.ydbio.2011.11.012

Maumus, F., Allen, A. E., Mhiri, C., Hu, H., Jabbari, K., Vardi, A., et al. (2009). Potential impact of stress activated retrotransposons on genome evolution in a marine diatom. *BMC Genomics*, *10*, 624. http://doi.org/10.1186/1471-2164-10-624

McLaughlin, S. (2017). Evidence that polycystins are involved in Hydra cnidocyte discharge. *Invertebrate Neuroscience : in*, *17*(1), 1. http://doi.org/10.1007/s10158-016-0194-3

Miller, D. J., Hemmrich, G., Ball, E. E., Hayward, D. C., Khalturin, K., Funayama, N., et al. (2007). The innate immune repertoire in cnidaria--ancestral complexity and stochastic gene loss. *Genome Biology*, *8*(4), R59. http://doi.org/10.1186/gb-2007-8-4-r59

Myant, K., Termanis, A., Sundaram, A. Y. M., Boe, T., Li, C., Merusi, C., et al. (2011). LSH and G9a/GLP complex are required for developmentally programmed DNA methylation. *Genome Research*, *21*(1), 83–94. http://doi.org/10.1101/gr.108498.110

Nevalainen, T. J., Peuravuori, H. J., Quinn, R. J., Llewellyn, L. E., Benzie, J. A. H., Fenner, P. J., & Winkel, K. D. (2004). Phospholipase A2 in cnidaria. *Comparative Biochemistry and Physiology. Part B, Biochemistry & Molecular Biology*, *139*(4), 731–735. http://doi.org/10.1016/j.cbpc.2004.09.006

Nicholson, D. W., & Thornberry, N. A. (1997). Caspases: killer proteases. *Trends in Biochemical Sciences*, *22*(8), 299–306.

Opipari, A. W., Boguski, M. S., & Dixit, V. M. (1990). The A20 cDNA induced by tumor necrosis factor alpha encodes a novel type of zinc finger protein. *The Journal of Biological Chemistry*, *265*(25), 14705–14708.

Peres, R., Reitzel, A. M., Passamaneck, Y., Afeche, S. C., Cipolla-Neto, J., Marques, A. C., & Martindale, M. Q. (2014). Developmental and light-entrained expression of melatonin and its relationship to the circadian clock in the sea anemone Nematostella vectensis. *EvoDevo*, *5*, 26. http://doi.org/10.1186/2041-9139-5-26

Porté, S., Valencia, E., Yakovtseva, E. A., Borràs, E., Shafqat, N., Debreczeny, J. E., et al. (2009). Three-dimensional structure and enzymatic function of proapoptotic human p53-inducible quinone oxidoreductase PIG3. *The Journal of Biological Chemistry*, *284*(25), 17194–17205. http://doi.org/10.1074/jbc.M109.001800

Pörtner, H. O. (2002). Climate variations and the physiological basis of temperature dependent biogeography: systemic to molecular hierarchy of thermal tolerance in animals. *Comparative Biochemistry and Physiology. Part a, Molecular & Integrative Physiology*, *132*(4), 739–761.

Pratlong, M., Haguenauer, A., Chabrol, O., Klopp, C., Pontarotti, P., & Aurelle, D. (2015). The red coral (Corallium rubrum) transcriptome: a new resource for population genetics and local adaptation studies. *Molecular Ecology Resources*, *15*(5), 1205–1215. http://doi.org/10.1111/1755-0998.12383

Ramos-Silva, P., Kaandorp, J., Huisman, L., Marie, B., Zanella-Cléon, I., Guichard, N., et al. (2013). The skeletal proteome of the coral Acropora millepora: the evolution of calcification by co-option and domain shuffling. *Molecular Biology and Evolution*, *30*(9), 2099–2112. http://doi.org/10.1093/molbev/mst109

Reitzel, A. M., Tarrant, A. M., & Levy, O. (2013). Circadian clocks in the cnidaria: environmental entrainment, molecular regulation, and organismal outputs. *Integrative and Comparative Biology*, *53*(1), 118–130. http://doi.org/10.1093/icb/ict024

Reynolds, W. S., Schwarz, J. A., & Weis, V. M. (2000). Symbiosis-enhanced gene expression in cnidarian-algal associations: cloning and characterization of a cDNA, sym32, encoding a possible cell adhesion protein. *Comparative Biochemistry and Physiology. Part a, Molecular & Integrative Physiology*, *126*(1), 33–44.

Rivest, E. B., & Hofmann, G. E. (2014). Responses of the Metabolism of the Larvae of Pocillopora damicornis to Ocean Acidification and Warming. *PloS One*, *9*(4), e96172. http://doi.org/10.1371/journal.pone.0096172.s004

Schallreuter, K. U., Gibbons, N. C. J., Zothner, C., Abou Elloof, M. M., & Wood, J. M. (2007). Hydrogen peroxide-mediated oxidative stress disrupts calcium binding on calmodulin: more evidence for oxidative stress in vitiligo. *Biochemical and Biophysical Research Communications*, *360*(1), 70–75. http://doi.org/10.1016/j.bbrc.2007.05.218

Schwarz, J. A., & Weis, V. M. (2003). Localization of a symbiosis-related protein, Sym32, in the Anthopleura elegantissima-Symbiodinium muscatinei Association. *The Biological Bulletin*, *205*(3), 339–350. http://doi.org/10.2307/1543297

Sterner, D. E., & Berger, S. L. (2000). Acetylation of histones and transcription-related factors. *Microbiology and Molecular Biology Reviews*, *64*(2), 435–459.

Tashian, R. E. (1989). The carbonic anhydrases: widening perspectives on their evolution, expression and function. *BioEssays*, *10*(6), 186–192. http://doi.org/10.1002/bies.950100603

Vervoort, M., Meulemeester, D., Béhague, J., & Kerner, P. (2016). Evolution of Prdm Genes in Animals: Insights from Comparative Genomics. *Molecular Biology and Evolution*, *33*(3), 679–696. http://doi.org/10.1093/molbev/msv260

Vidal-Dupiol, J., Vidal-Dupiol, J., Adjeroud, M., Adjeroud, M., Roger, E., Roger, E., et al. (2009). Coral bleaching under thermal stress: putative involvement of host/symbiont recognition mechanisms. *BMC Physiology*, *9*, 14. http://doi.org/10.1186/1472-6793-9-14

Vize, P. D. (2009). Transcriptome analysis of the circadian regulatory network in the coral Acropora millepora. *The Biological Bulletin*, *216*(2), 131–137. http://doi.org/10.1086/BBLv216n2p131

Voolstra, C. R., Schnetzer, J., Peshkin, L., Randall, C. J., Szmant, A. M., & Medina, M. (2009). Effects of temperature on gene expression in embryos of the coral Montastraea faveolata. *BMC Genomics*, *10*, 627. http://doi.org/10.1186/1471-2164-10-627

Vulpe, C. D., Kuo, Y. M., Murphy, T. L., Cowley, L., Askwith, C., Libina, N., et al. (1999). Hephaestin, a ceruloplasmin homologue implicated in intestinal iron transport, is defective in the sla mouse. *Nature Genetics*, *21*(2), 195–199. http://doi.org/10.1038/5979

Weis, V. M. (2008). Cellular mechanisms of Cnidarian bleaching: stress causes the collapse of symbiosis. *The Journal of Experimental Biology*, *211*(Pt 19), 3059–3066. http://doi.org/10.1242/jeb.009597

Wessler, S. R. (1996). Turned on by stress. Plant retrotransposons. *Current Biology : CB*, *6*(8), 959–961.

Wood-Charlson, E. M., Hollingsworth, L. L., Krupp, D. A., & Weis, V. M. (2006). Lectin/glycan interactions play a role in recognition in a coral/dinoflagellate symbiosis. *Cellular Microbiology*, *8*(12), 1985–1993. http://doi.org/10.1111/j.1462-5822.2006.00765.x

Yakovleva, I. M., Baird, A. H., Yamamoto, H. H., Bhagooli, R., Nonaka, M., & Hidaka, M. (2009). Algal symbionts increase oxidative damage and death in coral larvae at high temperatures.

Yoda, H., Hiroi, Y., & Sano, H. (2006). Polyamine oxidase is one of the key elements for oxidative burst to induce programmed cell death in tobacco cultured cells. *Plant Physiology*, *142*(1), 193–206. http://doi.org/10.1104/pp.106.080515
